# Sex specific disruptions in Protein Kinase Cγ signaling in a mouse model of Spinocerebellar Ataxia Type 14

**DOI:** 10.1101/2025.02.10.637267

**Authors:** Sarah A. Wolfe, Yuliang Ma, Caila A. Pilo, Carly Chang, Majid Ghassemian, Amanda J. Roberts, Sang Ryeul Lee, George Gorrie, Susan S. Taylor, Alexandra C. Newton

## Abstract

Spinocerebellar Ataxia Type 14 (SCA14) is an autosomal dominant neurodegenerative disease caused by mutations in the gene encoding protein kinase C gamma (PKCγ), a Ca^2+^/diacylglycerol (DG)-dependent serine/threonine kinase dominantly expressed in cerebellar Purkinje cells. These mutations impair autoinhibitory constraints to increase the basal activity of the kinase, resulting in deficits in the cerebellum that are not observed upon simple deletion of the gene, and severe ataxia. To better understand the phenotypic impact of aberrant PKCγ signaling in disease pathology, we developed a knock-in murine model of the SCA14 mutation ΔF48 in PKCγ. This fully penetrant mutation is severe in humans and is mechanistically informative as it has high basal activity but is unresponsive to agonist stimulation. Genetic, behavioral, and molecular testing revealed that ΔF48 PKCγ SCA14 mice have ataxia related phenotypes and an altered cerebellar phosphoproteome, effects that are more severe in male mice. Analysis of existing human data reveal that SCA14 has a significantly earlier age of onset for males compared with females. Our data from this clinically relevant mutation suggest that enhanced basal activity of PKCγ is necessary and sufficient to cause ataxia and that treatment strategies to modulate aberrant PKCγ may be particularly beneficial in males.

**Summary:** New mouse model of Spinocerebellar Ataxia Type 14 containing a clinically relevant mutation in PKCγ identified underlying drivers of the disease and neuroprotection in females.

## Introduction

The neurodegenerative disease Spinocerebellar Ataxia Type 14 (SCA14) is caused by missense mutations in the Ca^2+^/diacylglycerol (DG)-dependent Ser/Thr kinase, protein kinase C gamma (PKCγ)(1). SCAs are a group of autosomal dominant diseases characterized by cerebellar dysfunction, progressive ataxia, loss of motor coordination, increased disease severity with age, and cognitive-affective disturbances(2-15). SCAs are genetically heterogeneous, with close to 50 genetic causes for the disease identified(16, 17). Germline missense mutations in PKCγ were first identified two decades ago and there are now over 75 known mutations PKCγ that define SCA14(1, 2, 18). The majority of SCAs are driven by dysregulated Ca^2+^ homeostasis, including mutations in genes encoding the primary inositol 1,4,5-trisphosphate (IP_3_) receptor in the brain (SCA15, 16, 29), Ca^2+^ channel subunits (SCA25, 26), and voltage-gated K+ channels that impact voltage-gated Ca^2+^ channel function (SCA13, 19, 22). As a key transducer of Ca^2+^ signals, aberrant PKCγ signaling is associated with most SCAs(19-22). Understanding the pathology of PKCγ mutations in SCA14 may open avenues for therapeutically targeting SCAs in general, particularly given the druggability of protein kinases.

PKC isozymes play key roles in normal brain physiology by regulating neuronal functions such as synapse morphology, receptor turnover, and cytoskeletal integrity. As with all conventional PKC isozymes, PKCγ is tightly regulated and is maintained in an ‘off’ autoinhibited state by a set of N-terminal regulatory domains that prevent the C-terminal kinase domain from phosphorylating its substrates(21-24). Notably, the substrate-binding site of PKCγ binds an autoinhibitory pseudosubstrate segment that maintains PKCγ in an inactive conformation; binding to second messengers Ca^2+^ and DG releases the pseudosubstrate, allowing PKC to adopt the ‘on’ and signaling-competent conformation. These second messengers are produced by phospholipase C (PLC)-catalyzed hydrolysis of phosphatidylinositol 4,5-bisphosphate (PIP_2_) to produce IP_3_, which induces intracellular Ca^2+^ release, and the allosteric activator DG. Binding of Ca^2+^ to the C2 domain recruits PKCγ to the plasma membrane, where it binds the membrane-embedded DG via the C1B domain, to relieve autoinhibition and permit efficient substrate phosphorylation(21-27). Metabolism of DG and reduction of cytosolic Ca^2+^ cause PKCγ to re-autoinhibit, resulting in re-autoinhibition and the ‘off’ state(28, 29). Improperly autoinhibited PKC is degraded by a quality control mechanism that ensures that aberrantly active PKC does not accumulate in the cell(30, 31).

SCA14 mutations impair autoinhibitory contacts to increase basal activity of PKCγ, resulting in ‘leaky’ signaling(21, 32-34). However, many of these mutations uniquely evade the quality control degradation mechanism that can be induced by the DG functional analogue, phorbol esters, which is not metabolized and traps PKC in the ‘on’ conformation(21, 32). One of the most severe variants, deletion of Phe48 (ΔF48) in PKCγ, is in the C1A domain immediately following the pseudosubstrate. ΔF48 PKCγ not only has reduced autoinhibition resulting in higher basal activity but also has uncoupled communication between the pseudosubstrate and the kinase domain such that the enzyme is trapped in an unresponsive but leaky signaling state. The unique unresponsiveness of this variant to DG/Ca^2+^-induced activation suggests that the unregulated leaky basal signaling of this SCA14 variant is necessary and sufficient to drive the pathology(21, 32, 33).

PKCγ is dominantly expressed in Purkinje cells(34-43), a major cell type in the cerebellum. Purkinje cells are GABAergic neurons essential to cerebellar signaling and function. They form highly branched dendritic trees and are innervated throughout the cerebellar layers [molecular layer (ML), Purkinje cell layer (PCL), and granular layer (GL)](44-47). PKCγ signaling plays an important role in Purkinje cell function, formation, and innervation in the cerebellum, and disruptions in proper Purkinje cell development occur when PKC activity is dysregulated(34-43, 48-53). Other PKC isoforms are also present in Purkinje cells including α, δ, and η(36, 54, 55). SCA pathogenesis can be driven by progressive neurodegeneration of Purkinje cells, or by dysfunction and altered firing patterns of Purkinje cells that disrupt cerebellar signaling causing motor dysfunction(6, 7, 34, 48-52, 56, 57, 58). It is noteworthy that PKCγ knock-out mice lack evident neurodegeneration, revealing that it is not the absence of PKCγ that drives neurodegeneration in SCA14, but the presence of basally active PKCγ and aberrant signaling(34, 41, 53, 59, 60). How this dysregulated PKCγ signaling drives the pathology of SCA14 remains to be elucidated.

Here we have generated a mouse model of SCA14 containing the severe mutation, ΔF48 PKCγ, to better understand how basally active PKCγ drives the pathology of SCA14. The ΔF48 PKCγ mouse displayed ataxia-associated phenotypes that were more severe in males compared with females. These mice displayed significantly reduced PKCγ protein levels with no reduction in PKC substrate phosphorylation, revealing that the constitutive basal activity of ΔF48 PKCγ compensated for reduced PKC levels. Immunohistochemical analyses revealed altered cerebellar Purkinje cell arborization, but a lack of overt neurodegeneration. Phosphoproteomic analysis revealed significant alterations in cerebellum of the ΔF48 PKCγ mice compared to wild-type (WT) that primarily impacted neuronal and cytoskeletal pathways, consistent with deficits in Purkinje cells. Taken together with existing human data showing an earlier age of onset of SCA14 symptoms in male compared with female patients, our data from the mouse model suggest that this disease is more severe in males compared with females.

## Results

### ΔF48 PKCγ causes sex-specific ataxia related deficits in motor functions in a mouse model of SCA14

ΔF48 PKCγ has enhanced basal activity due to impaired autoinhibition, increased turnover due to quality control degradation, resistance to phorbol ester-induced degradation (which requires an intact C1A domain), and insensitivity to DG/Ca^2+^-induced activation(21, 32). These severe biochemical defects of ΔF48 PKCγ are associated with early age of onset and high disease severity(11, 32). To understand the pathology of ΔF48 PKCγ-induced SCA14, a ΔF48 PKCγ knock-in mouse was generated with CRISPR/Cas9-mediated genome editing methods.

To assess the viability of mice with SCA14 genotypes [heterozygous WT/ΔF48 (HET) or homozygous ΔF48/ΔF48 (HOM)], the birth rates of each genotype were compared from 183 offspring born from HET crosses. Female mice had expected Mendelian inheritance ratios (WT: 29%, HET: 46%, HOM: 25%), whereas male mice had fewer than expected HOM mice born (WT: 30%, HET: 53%, HOM: 17%). In HOM offspring, male (33%) and female (67%) birth ratios were significantly different from expected ratios (**Figure 1a**). This suggests that male HOM mice have reduced birth rates and possible developmental deficits resulting from aberrant PKCγ signaling(2-8, 15, 41). A hallmark of SCA14 is cerebellar dysfunction leading to progressive ataxia and loss of motor function(2-15). To evaluate motor function in SCA14 mice with the ΔF48 PKCγ mutation, behavioral tests used to measure motor function including a wire hang test, a rotarod test, a treadmill walking test, a ladder rung test, and a grip strength test were performed on adult mice over 5 months old. Using the wire hang test, grip and muscle strength were assessed by measuring latency to fall from a wire. HOM mice had a markedly reduced latency to fall time by ∼70% in both sexes compared to WT mice, and a significantly reduced latency to fall time by 70% in females and 54% in males compared to HET mice. Male HET mice also had a significant 32% reduction in hang time compared to WT, whereas female HET mice performed equivalently to WT (**Figure 1b**). The rotarod test was used to evaluate locomotor coordination and balance by measuring latency to fall from a rotating rod. Both female and male HOM mice had significantly decreased latency to fall times compared to WT (reduced by 45% in females and 64% in males) and HET mice (reduced by 55% in females and 62% in males) (**Figure 1c**). To determine deficits in walking, SCA14 mice were evaluated with a treadmill task in which the mouse’s percent time spent walking ahead of the bumper/rear of a treadmill was measured. Only male HOM mice were found to have a significantly reduced time spent walking compared to WT by 23% (**Figure 1d**). The ladder rung test was used to measure skilled walking and coordination by measuring paw slips as mice walk along a horizontal ladder. No significant differences were detected with this test, however, a strong trend towards an increase in the number of slips measured in the male and female HOM mice was observed (**Figure 1e**). No significant differences were observed in forelimb grip strength between genotypes, however, a trend towards a decrease in grip strength was observed in male HET and HOM mice (∼10% reduced) (**Figure 1f**). These behavioral tests indicate that SCA14 mice have ataxia-associated phenotypes that are more severe in males.

**Figure 1:**
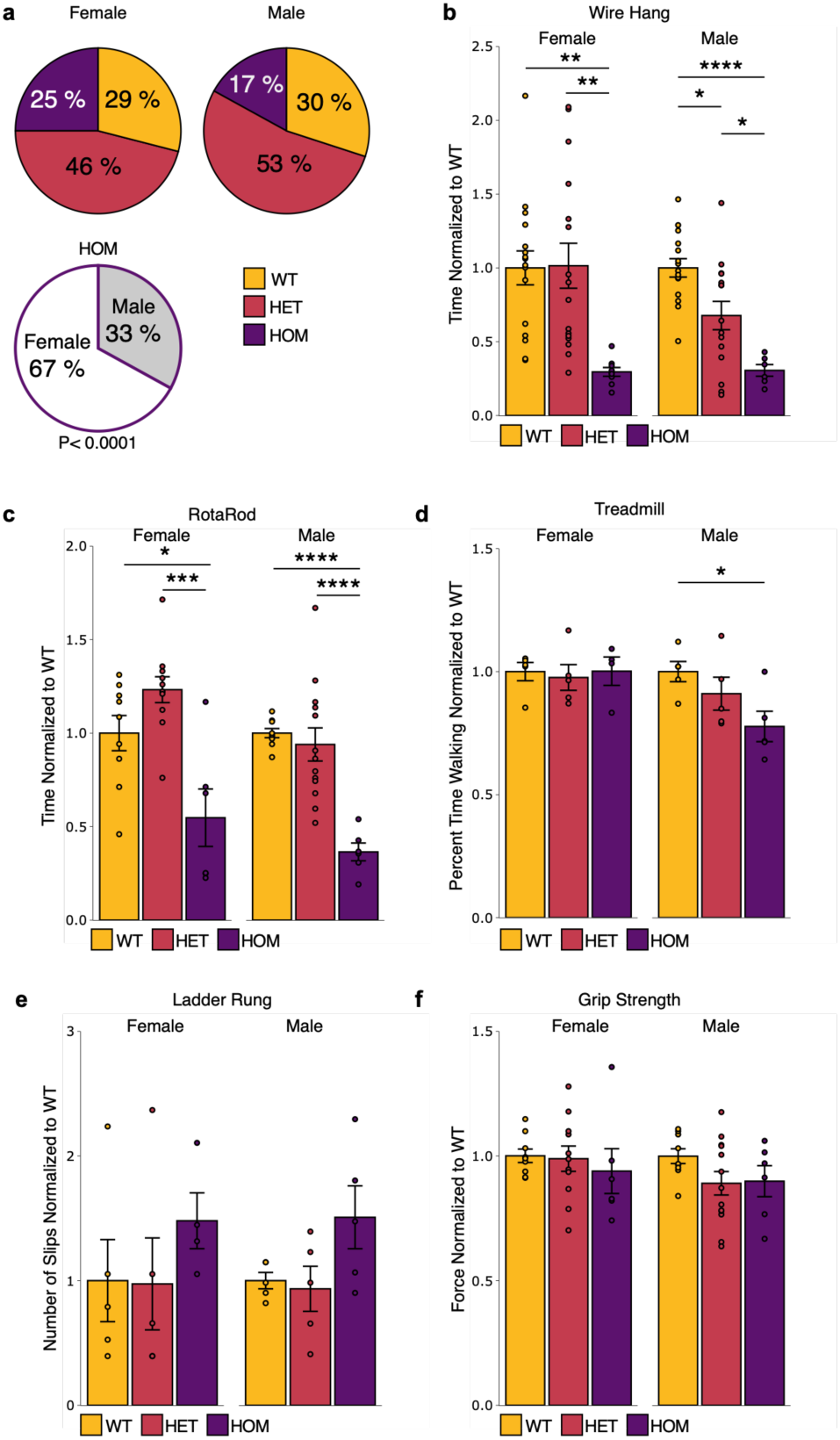
ΔF48 PKCγ causes ataxia-associated deficits in motor function. Viability of SCA14 mice containing the ΔF48 PKCγ mutation were determined by evaluating offspring born from HET crosses. **a)** Pie charts show genetic inheritance ratios for WT (yellow), HET (red), and HOM (purple) female and male offspring. Significance was determined by χ^2^for expected ratios (p<0.0001, N=182 mice). SCA14 mice were evaluated for ataxic-related behaviors using a battery of motor function test in all genotypes (WT, yellow; HET, red; HOM, purple) and sexes including **b)** Wire Hang test [Female (One-way ANOVA: F(2,38)=7.7, p=0.0016; Tukey post-hoc test: WT vs HET p=0.99, WT vs HOM p=0.0032, HET vs HOM p=0.0027), Males (One-way ANOVA: F(2,33)=13, p=0.000069; Tukey post-hoc test: WT vs HET p=0.013, WT vs HOM p=0.000065, HET vs HOM p=0.033)], **c)** Rotarod test [Female (One-way ANOVA, F(2,23)=11, p=0.00040; Tukey post-hoc test, WT vs HET p=0.19, WT vs HOM p=0.017, HET vs HOM p=0.00026), Male (One-way ANOVA, F(2,25)=16, p=0.000033; Tukey post-hoc test, WT vs HET p=0.82, WT vs HOM p=0.000062, HET vs HOM p=0.000097)], **d)** Treadmill test [Female (One-way ANOVA, F(2,11)=0.088, p=0.92), Male (One-way ANOVA, F(2,12)=3.8, p=0.052; Tukey post-hoc test, WT vs HET p=0.53, WT vs HOM p=0.044, HET vs HOM p=0.27)], **e)** Ladder Rung test, and **f)** Grip Strength test. Data is normalized to WT where multiple cohorts were combined from mice (N=5-16 mice per group). Bar graphs represented quantification of mean±S.E.M. Significance was determined by One-way ANOVA with Tukey’s post hoc (p= *<0.05, **<0.01, ***<0.001, ***<0.0001).

### SCA14 mice have altered Purkinje cell morphology

The cerebellum is a key mediator of motor function, and cerebellar Purkinje cell neurodegeneration and/or dysfunction is a dominant feature of ataxia(34, 44-46, 48, 57, 58). To determine if adult SCA14 mice with the ΔF48 PKCγ mutation have deficits in cerebellar morphology indicative of ataxic pathology, we assessed the Purkinje cells of the cerebellum using immunofluorescence and Western blot analysis in all genotypes (WT, HET, HOM) and sexes using the Purkinje cell specific marker Calbindin D28k. Western blot analysis identified no alterations in Calbindin D28k protein abundance in cerebellar homogenate from adult SCA14 mice compared to WT in either male or female mice (**Figure 2a**), indicative of a lack of overt Purkinje cells neurodegeneration. To further assess the Purkinje cells, immunofluorescent staining of sagittal cerebellar slices (depicted and labeled in **Figure 2b**) was performed. Overall cerebellar morphology and Calbindin D28k intensity were visually similar in WT and SCA14 mice of both sexes (**Figure 2c**); however, upon closer inspection of an enlarged image of the Purkinje cell layer and molecular layer (indicated as the yellow box in Figure 2b), variations in Purkinje cells became more apparent. A ΔF48 PKCγ gene dose-dependent reduction (WT > HET > HOM) in dendritic arborization was observed and was particularly evident in male mice, as visualized by a reduction in Purkinje cell dendrite length and Calbindin D28k staining in the molecular layer **(Figure 2d)**. To measure potential differences, we selected the cerebellar layers where Purkinje cells are present (Purkinje cell layer and molecular layer) through all lobules of the cerebellum and analyzed the intensity of Calbindin D28k staining, the density of Purkinje cells, and thickness of the layers to assess Purkinje cell arborization. First, the mean intensity of Calbindin D28k staining was quantified, and no differences were observed in female ΔF48 PKCγ mice compared to WT; however, male ΔF48 PKCγ mice had reduced Calbindin D28k levels compared to WT, with a significant 33% reduction in HET males compared to WT males (**Figure 2e**). Next, the linear density of Purkinje cell somas was measured per distance along the Purkinje cell layer to identify neurodegeneration of Purkinje cells in SCA14 mice. Although, no significant loss of Purkinje cells was found, both male and female SCA14 genotypes trended down in Purkinje cell density (**Figure 2f**). Lastly, the thickness of the Purkinje cell layer and molecular layer was measured to assess Purkinje cell arborization. The average width of the Purkinje cell layer and molecular layer was measured, and a ΔF48 PKCγ gene dose-dependent reduction in thickness was identified in male mice with a significant average reduction of ∼23μm in HOM compared to WT males, indicating a reduction in dendritic length (**Figure 2g**). These data suggest that overt neurodegeneration or loss of Purkinje cells was not prevalent in the cerebellum of SCA14 mice, however, Purkinje cells had reduced dendritic branching and length, and this altered morphology was most apparent in the male mice.

**Figure 2:**
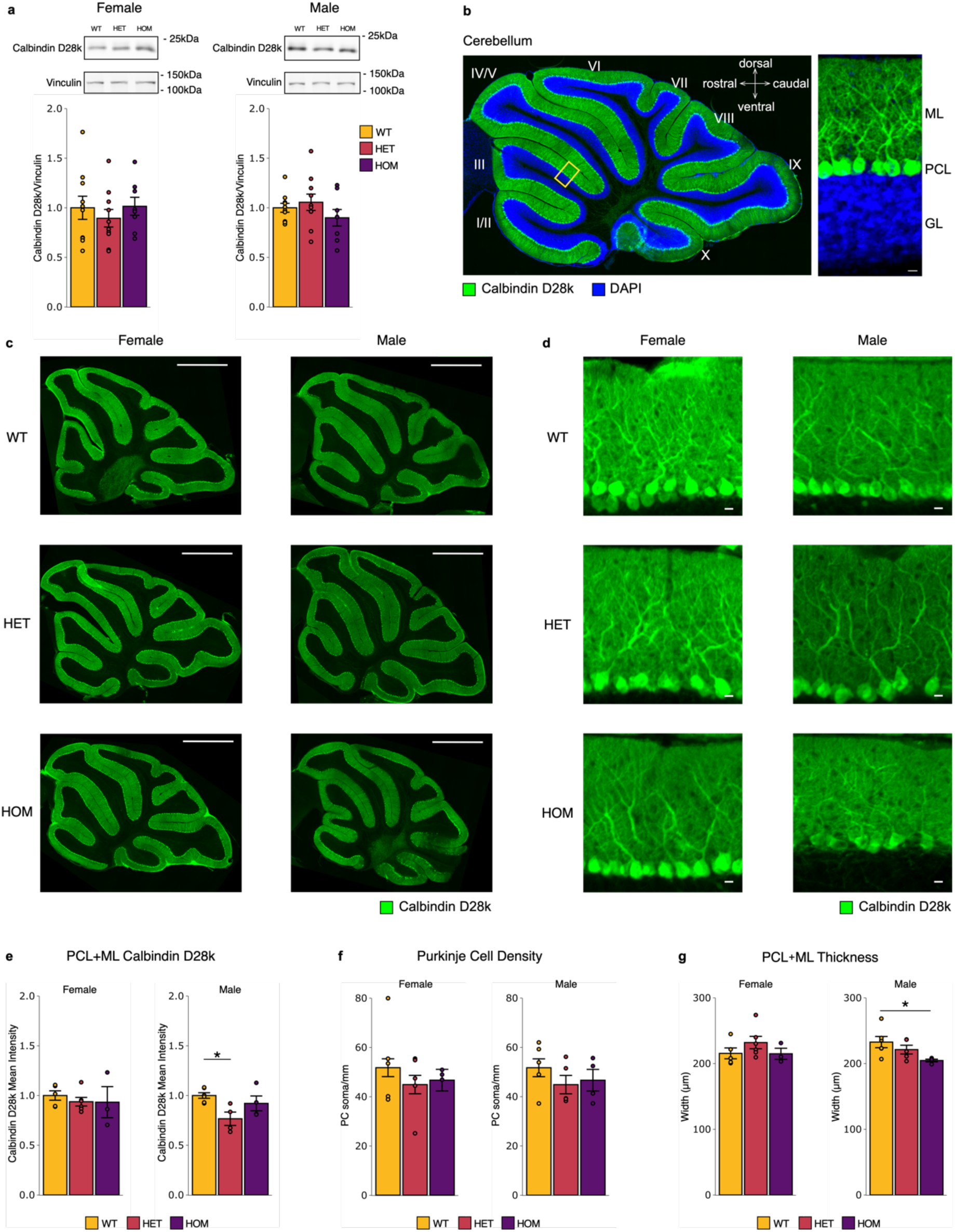
Cerebellum and Purkinje cell morphology are altered in SCA14 mice. The Purkinje cell marker, Calbindin D28k was used to assess neuronal morphology in SCA14 mice. **a)** Western blot analysis of whole cerebellar lysate from SCA14 mice of all genotypes (WT, yellow; HET, red; HOM, purple) and sexes was performed (N=8-10 mice per group). No significant differences were identified. Bar graphs represent quantification of mean±S.E.M. normalized to WT. Fluorescence immunohistochemistry of Calbindin D28k was performed on fixed sagittal brain sections from all genotypes (WT, HET, HOM) and sexes (N=3-6 mice). **b)** Example image indicates orientation of all cerebellum images, and yellow box indicates approximate location of magnified images, shown to the right with labeled Purkinje cell layer (PCL), molecular layer (ML), and granular layer (GL) (Calbindin D28k, green; DAPI, blue). Representative images show intensity and localization of Calbindin D28k (green) in **c)** whole cerebellum (Scale bar=1mm) and **d)** PCL and ML magnification (Scale bar=10μm). Quantification of PCL and ML of cerebellar images (N=3-6 mice) measured **e)** the intensity of Calbindin D28k labeling [Female (One-way ANOVA, F(2,11)=0.29, p=0.75; Tukey post-hoc test, WT vs HET p= 0.78, WT vs HOM p=0.82, HET vs HOM p=0.99), Male (One-way ANOVA, F(2,11)=5.0, p=0.028; Tukey post-hoc test, WT vs HET p=0.023, WT vs HOM p=0.55, HET vs HOM p=0.18)], **f)** the linear density of the Purkinje cells, and **g)** the width of the PCL and ML indicative of dendrite length [Female (One-way ANOVA, F(2,11)=1.2, p=0.34; Tukey post-hoc test, WT vs HET p=0.40, WT vs HOM p=1.0, HET vs HOM p=0.48), Male (One-way ANOVA, F(2,12)=3.6, p=0.058; Tukey post-hoc test, WT vs HET p=0.49, WT vs HOM p=0.047, HET vs HOM p=0.31)]. Bar graphs represented quantification of mean±S.E.M. Significance was determined by One-way ANOVA with Tukey’s post hoc (p= *<0.05, **<0.01, ***<0.001, ***<0.0001).

### SCA14 mice have reduced total PKC levels in the cerebellum

Reports indicate that PKCγ protein expression is reduced in SCA14, including both human and mouse model data(2, 34, 53, 56). To determine whether ΔF48 PKCγ affected PKC protein expression in the cerebellum, immunofluorescent staining of cerebellar sagittal slices and Western blot analysis of whole cerebellar homogenate was performed on all genotypes (WT, HET, HOM) and sexes for PKCγ and PKCα, another conventional PKC highly expressed in Purkinje cells(36, 54, 55). Western blot analysis of cerebellar tissue from SCA14 mice identified a ΔF48 PKCγ gene dose-dependent decrease in total PKCγ in the cerebellum. Analysis of HET mice showed a decrease in PKCγ by 41% in females and 27% in males, and HOM mice had severely reduced PKCγ levels by ∼90% in both sexes compared to WT. HOM mice also had significantly less PKCγ levels compared to HET mice in both sexes (83% less in females and 87% less in males) (**Figure 3a**). This loss in PKCγ was observed throughout all brain regions that express PKCγ as observed in stained sagittal slices of whole brain (**Supplemental Figure 1**). To determine where this reduction occurred specifically in the cerebellum, immunofluorescent staining was performed for PKCγ and Calbindin D28k. As expected, PKCγ was localized to Purkinje cells (**Supplemental Figure 2**) and a similar reduction in PKCγ levels was observed in SCA14 mice of both sexes (**Figure 3b**). Quantification of the mean intensity of the PKCγ signal in the Purkinje cell layer and molecular layer region revealed a ∼25% decrease in PKCγ in HET mice for both sexes, and HOM mice displayed a decrease of 69% in females and 62% in males compared to WT (**Figure 3c**). This reduction in steady-state protein was not a result of altered transcription as RT-qPCR revealed no significant differences in PKCγ RNA expression in WT vs HOM mice (**Supplemental Figure 3**). Levels of PKCα were modestly reduced in the SCA14 mice. Western blot analysis of the cerebellum indicated that PKCα levels trended towards a decrease in SCA14 female mice and was significantly reduced by 26% in HOM male mice compared to WT male mice (**Figure 3d).** Slices Stained for PKCα (**Figure 3e; Supplemental Figure 2**) with quantification of PKCα intensity in the molecular layer and Purkinje cell layer had no significant changes in PKCα levels (**Figure 3f**). These results indicate that PKCγ is gene dose-dependently reduced in SCA14 mice, due to the increased turnover rate previously identified for ΔF48 PKCγ, and that PKCα is not compensating for a loss of PKCγ protein in the Purkinje cells.

**Figure 3:**
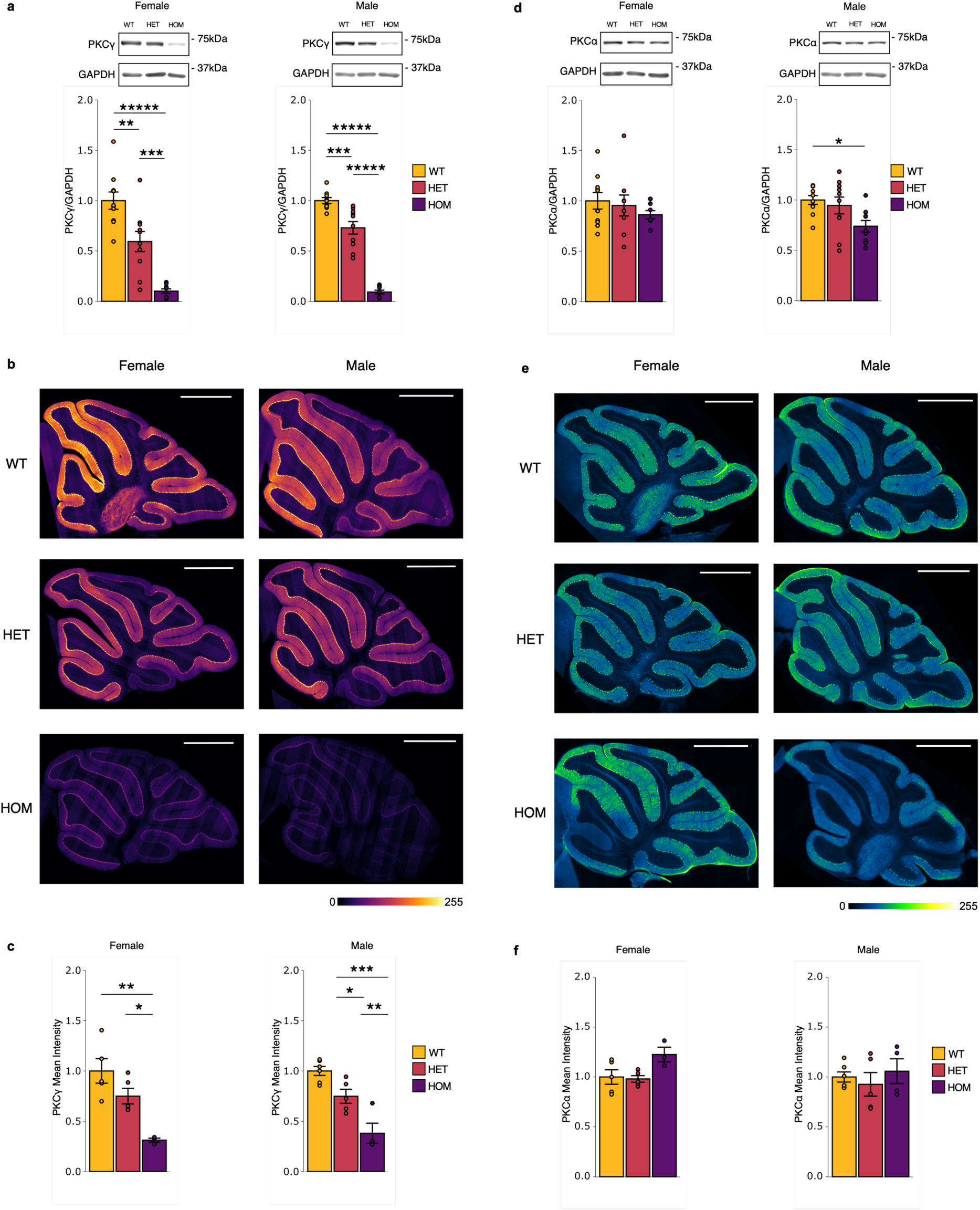
PKCγ and PKCα are gene dose dependently reduced in SCA14 mice. Western blot analysis of whole cerebellar homogenate from all genotypes (WT, yellow; HET, red; HOM, purple) and sexes (N=8-10 mice per group) was performed to assess cerebellar levels of **a)** PKCγ [Female (One-way ANOVA, F(2,26)=31.14, p=1.26e-07; Tukey post-hoc test, WT vs HET p=0.0031, WT vs HOM p<0.00001, HET vs HOM p=0.00057), Male (One-way ANOVA, F(2,24)=101.20, p=2.02e-12; Tukey post-hoc test, WT vs HET p=0.00057, WT vs HOM p<0.00001, HET vs HOM p<0.00001)], and **d)** PKCα [Female (One-way ANOVA, F(2,25)=0.76, p=0.48), Male (One-way ANOVA, F(2,26)=4.4, p=0.022; Tukey post-hoc test, WT vs HET p=0.82, WT vs HOM p=0.023, HET vs HOM p=0.081)]. Fluorescence immunohistochemistry was performed on fixed sagittal brain sections from all genotypes (WT, HET, HOM) and sexes (N=3-6 mice). Representative images show intensity and localization of **b)** PKCγ in whole cerebellum (Scale bar=1mm, respective color scale indicates relative intensity). Quantification of the PCL and ML (N=3-6 mice) measured the intensity of **c)** PKCγ localized to the PCL and ML [Female (One-way ANOVA, F(2,10)=11, p=0.0033; Tukey post-hoc test, WT vs HET p=0.18, WT vs HOM p=0.0025, HET vs HOM p= 0.036), Males (One-way ANOVA, F(2,12)=20, p=0.00015; Tukey post-hoc test, WT vs HET p=0.044, WT vs HOM p=0.00010, HET vs HOM p=0.0092)]. Representative images show intensity and localization of **e)** PKCα in whole cerebellum (Scale bar=1mm, respective color scale indicates relative intensity). Quantification of the PCL and ML (N=3-6 mice) measured the intensity of **f)** PKCα localized to the PCL and ML. Bar graphs are mean±S.E.M. normalized to WT. Significance was determined by One-way ANOVA with Tukey’s post hoc (p=*<0.05, **<0.01, ***<0.001, ***<0.0001).

### Total PKC substrate phosphorylation and other PKC isoforms are unaltered in SCA14 mice

Although SCA14 mice have significantly reduced levels of PKCγ protein, we reasoned that the enhanced basal activity of ΔF48 PKCγ could drive aberrant phosphorylation. To identify if the observed loss of PKCγ in the cerebellum of SCA14 mice affected overall PKC signaling, we measured total PKC substrate phosphorylation using Western blot analysis of cerebellar homogenate from all genotypes (WT, HET, HOM) and sexes. No significant differences in substrate phosphorylation were identified despite reduced levels of PKCγ and PKCα in SCA14 mice compared to WT in either sex (**Figure 4a**). We also tested phosphorylation of the Glycogen Synthase Kinase-3 beta (GSK3β), a protein known to have increased phosphorylation with other overactive, ataxia-associated mutations in PKCγ(32), and contains a bonafide PKC consensus RxxS site on serine 9 (Ser^9^)(32, 61). No significant differences were identified in phosphorylation of GSK3β (Ser^9^) in SCA14 mice in either sex (**Figure 4b**). To address whether other PKC isoforms may affect total PKC substrate phosphorylation and compensate for the reduction in PKCγ protein, we measured the expression of other PKC isoforms known to be present in cerebellar Purkinje cells, PKCδ and PKCη(36, 54, 55). Western blot analysis of revealed no significant differences in the levels of PKCδ (**Figure 4c**) or PKCη (**Figure 4d**) between SCA14 genotypes compared to WT in either sex. Thus, although PKCγ is dramatically reduced in the cerebellum of SCA14 mice, its higher leaky basal activity results in no overall change in PKC substrate phosphorylation, including phosphorylation of the previously identified phosphorylation site on GSK3β that has been shown to increase in ataxia models of overactive mutant PKCγ(32, 61).

**Figure 4:**
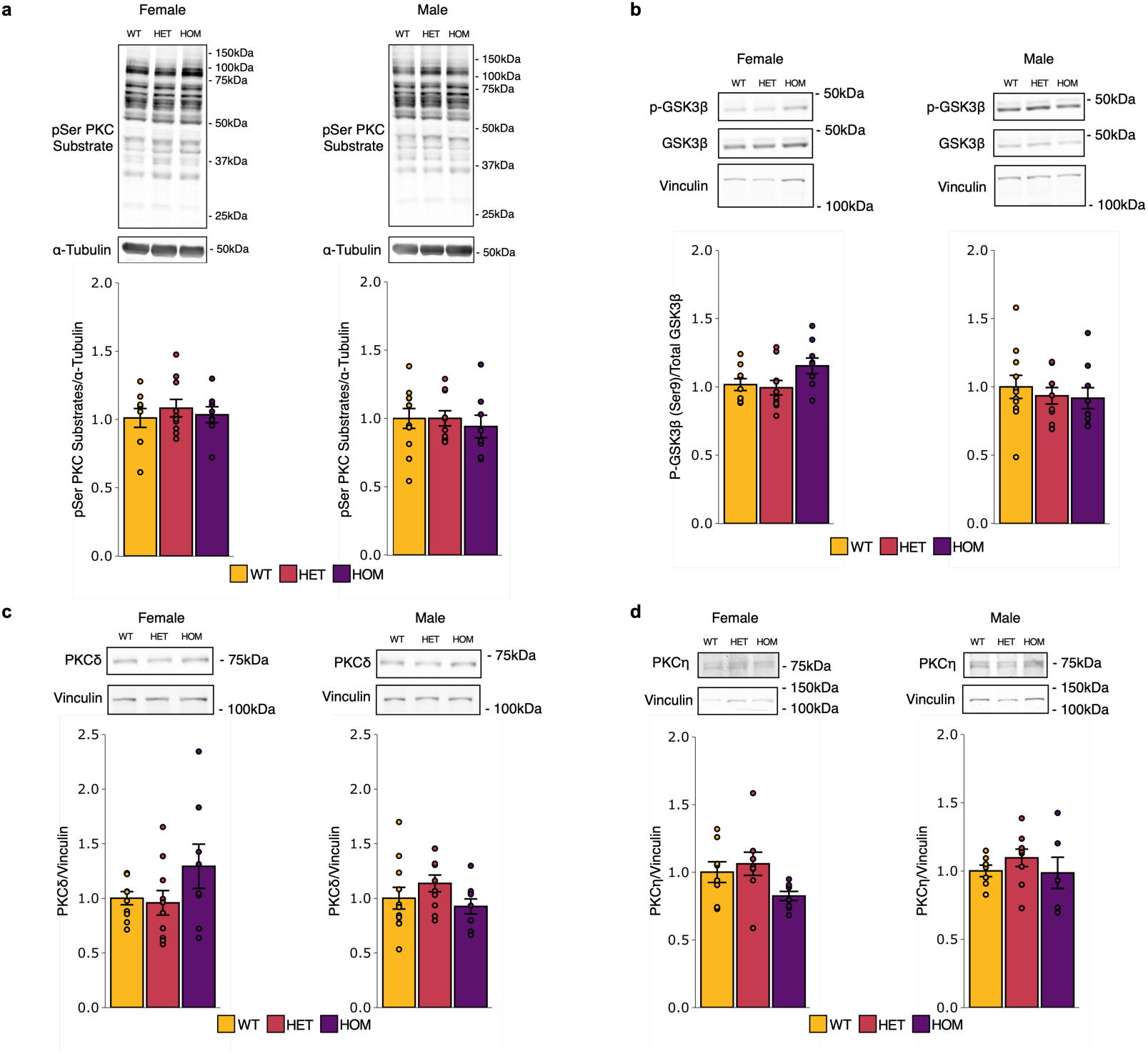
SCA14 mice have equivalent PKC substrate phosphorylation and PKC isozyme levels as WT mice. Western blot analysis was performed on whole cerebellar homogenate from all genotypes (WT, yellow; HET, red; HOM, purple) and sexes (N=6-11 mice per group) to assess pSer PKC substrate phosphorylation, GSK3β (Ser^9^) phosphorylation, and PKCδ and PKCη expression. Western blot analysis for **a)** pSer PKC substrate phosphorylation sites, **b)** phosphorylated over total GSK3β, **c)** PKCδ, and **d)** PKCη identified no significant differences between genotype or sex. Western blot data is normalized to WT. Bar graphs represented quantification of mean±S.E.M. Significance was determined by One-way ANOVA with Tukey’s post hoc.

### SCA14 mice have altered phosphoproteome

To identify the effects of the aberrant ΔF48 PKCγ signaling in the cerebellum, a phosphoproteomic analysis was carried out on protein extracted from whole cerebellar homogenate from all genotypes (WT, HET, HOM) and sexes. A total of 5216 quantifiable proteins and 7829 quantifiable phosphopeptides were detected. Phosphopeptide intensity was normalized to the corresponding protein intensity. Protein intensity was determined in a parallel proteome analysis, displayed in **Supplemental Figure 4**. Significant differences in phosphopeptides abundance (as determined by two sample t-test <0.05) were compared between genotypes in female and male mice. Despite there being no net increases in PKC substrate phosphorylation assessed by Western blot analysis (**Figure 4a**), alterations in specific phosphopeptides were detected in the phosphoproteomic analysis of the cerebellums (**Figure 5**). An upset plot was used to display the significant differentially abundant phosphopeptides found between each SCA14 genotype compared to WT (sets: WT vs HET and WT vs HOM) in male and female mice (**Figure 5a**). Male HOM mice had the greatest number of phosphopeptides that differed compared to WT and the greatest number of differentially abundant phosphopeptides not shared with any other sets (Set size: Female: WT vs HET= 246, HOM = 184; Male: WT vs HET= 212, WT vs HOM = 257). This is consistent with phenotypic data that indicated HOM male mice have the most severe defects. The intersections indicate that relatively few phosphopeptides were shared between sets, with the male HET and male HOM having the most in common at 37, followed by female HET and female HOM at 34 phosphopeptides (**Figure 5a**). No differentially abundant phosphopeptides were found in common between all sets. These data suggesting that ΔF48 PKCγ rewires the phosphoproteome sex-specifically.

**Figure 5:**
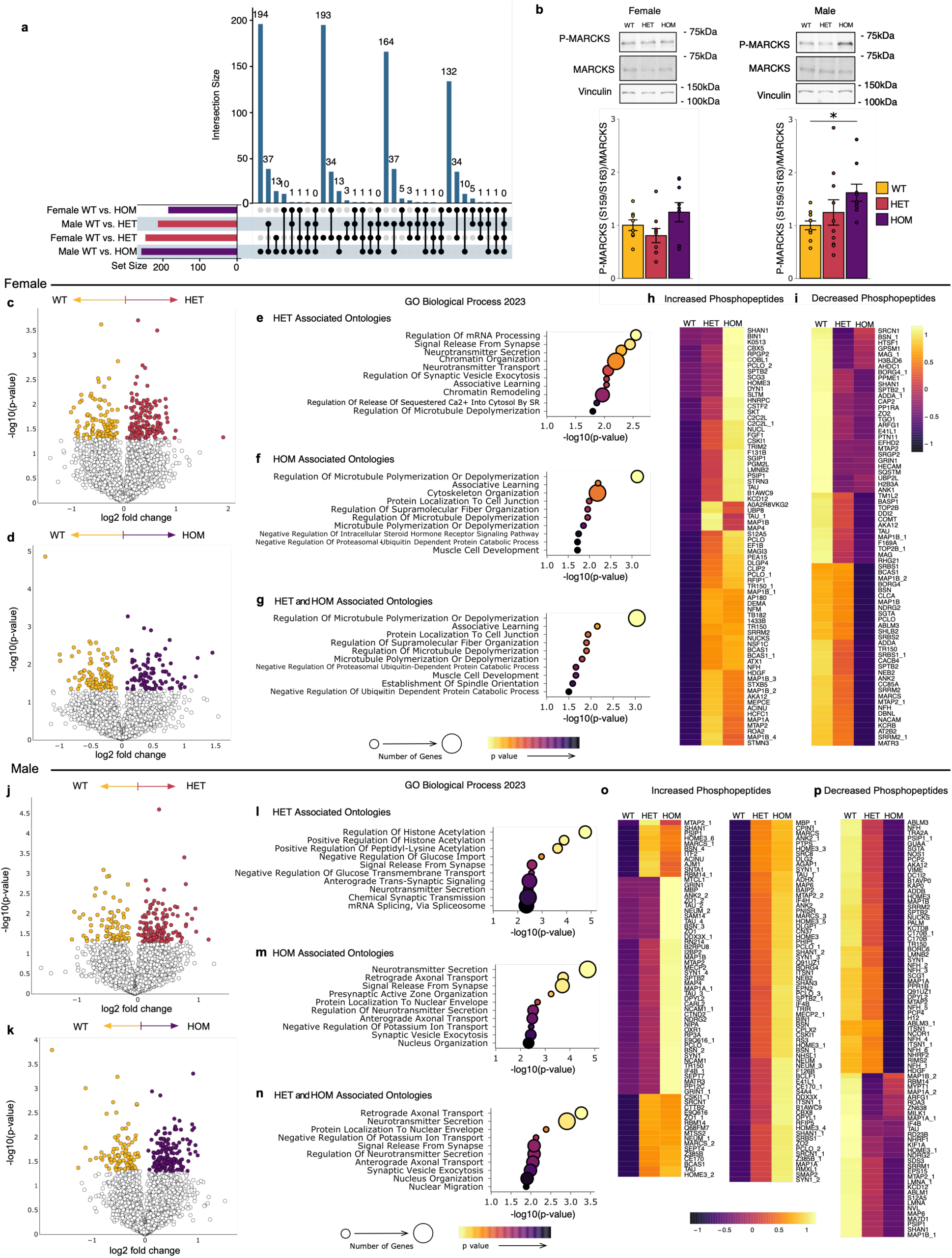
ΔF48 PKCγ rewires the cerebellar phosphoproteome in SCA14 mice. Phosphoproteomic analysis was carried out on protein extracted from whole cerebellar homogenate from all genotypes (WT, yellow; HET, red; HOM, purple) and sexes (N=3 mice per group). **a)** An upset plot summarizes the quantity of significantly different phosphopeptides and shared phosphopeptides identified between groups (p<0.05). **b)** Western blot analysis was performed on whole cerebellar homogenate (N=8-10 mice per group) to assess MARCKS phosphorylation (Ser^159^/Ser^163^). [Female (One-way ANOVA, F(2,23)=2.4, p=0.11; Tukey post-hoc test, WT vs HET p=0.62, WT vs HOM p=0.48, HET vs HOM p=0.094), Males (One-way ANOVA, F(2,26)=3.1, p=0.063; Tukey post-hoc test, WT vs HET p=0.58, WT vs HOM p=0.051, HET vs HOM p=0.31)]. Bar graphs represent mean±S.E.M. Significance determined by One-way ANOVA with Tukey’s post hoc. Volcano plots show log-transformed p-values vs the log-transformed fold change quantified per change in phosphopeptide abundance between WT compared to **c)** HET and **d)** HOM in female mice. Color represents phosphopeptides with p-value<0.05. Dot plots indicate the top 10 most significantly enriched for gene ontologies from the differentially abundant phosphopeptides identified in **e)** WT vs HET, **f)** WT vs HOM, and **g)** similarly altered phosphopeptides (defined as phosphopeptides significantly altered in HOM vs WT with a corresponding trend, or significant change, in HET vs WT) in female mice. Similarly altered phosphopeptides are shown in heatmaps by those that **h)** increase or **i)** decrease with increasing ΔF48 alleles in females. In male mice, volcano plots show change in phosphopeptide abundance between WT compared to **j)** HET and **k)** HOM. Color represents phosphopeptides with p-value < 0.05. Dot plots indicate the top 10 most significantly enriched for gene ontologies from the differentially abundant phosphopeptides identified in **l)** WT vs HET, **m)** WT vs HOM, and **n)** similarly altered phosphopeptides in males. Color scale indicates p-value, dot size indicates number of genes per ontology (p<0.05). Similarly altered phosphopeptides are shown in heatmaps by those that **o)** increase or **p)** decrease with increasing ΔF48 alleles in males. Dot plot color scale indicates p-value, dot size indicates number of genes per ontology (p<0.05). Heatmap color scale indicates normalized phosphopeptides intensity with values centered and scaled by row. Protein names accompanied by a number designate multiple phosphorylation sites for the same protein.

To validate sex-specific changes, Western blot analysis was performed on cerebellum from mice of all genotypes (WT, HET, HOM) and sexes for phosphorylated (Ser^159^/Ser^163^) and total myristoylated alanine-rich C-kinase substrate (MARCKS), a bonafide PKC substrate. A significant increase in phosphorylation of MARCKS was observed in HOM males, but not females, compared to WT (**Figure 5b**). A similar increase was observed in the phosphoproteomic analysis for the same phosphorylation site (Ser^163^) in SCA14 male mice. Thus, ΔF48 PKCγ has significantly larger effects on the phosphorylation of diverse substrates, including the well-characterized MARCKS substrate.

To better understand signaling pathways and cellular functions differentially altered by the ΔF48 PKCγ variant, significant differences in phosphopeptide abundance with SCA14 genotypes were first compared in each sex separately. In female mice, volcano plots indicate phosphopeptides that were significantly increased (more abundant in the SCA14 genotype) or decreased (more abundant in WT) in HET (**Figure 5c**) and HOM (**Figure 5d**) compared to WT female mice. Corresponding gene ontology analysis was performed for significantly different phosphopeptides found in HET and HOM compared to WT female mice for biological processes (**Figure 5e-f**), cellular components (**Supplemental Figure 5a-b**), and molecular functions (**Supplemental Figure 5c-d**). The top 10 most significantly enriched for ontologies for biological processes are shown as assessed using EnrichR(62-64) (p<0.05). Gene ontology analysis was also performed on significantly different phosphopeptides associated with both HET and HOM female mice. These similarly associated phosphopeptides were determined as phosphopeptides that significantly altered in the HOM vs WT comparison and had a corresponding trend or significant change in the same direction in the HET vs WT comparison (biological processes, **Figure 5g**; cellular components, **Supplemental Figure 5e;** molecular functions **Supplemental Figure 5f**). These similarly altered phosphopeptides were displayed in heatmaps as those that were increased (**Figure 5h**) or decreased (**Figure 5i**) in the SCA14 genotypes compared to WT in females. As with the female mice, volcano plots (**Figure 5j-k**), gene ontologies (**Figure 5l-n; Supplemental Figure 5g-l**), and heatmaps (**Figure 5o-p**) are displayed using the same analysis for the male mice. Concerning the similarly associated phosphopeptides, 78% of all the differentially abundant phosphopeptides in HOM female mice were similarly associated in HET female mice, and 85% of all the differentially abundant phosphopeptides in HOM male mice were similarly associated in HET male mice. This suggesting that both SCA14 genotypes have similar effects but may be present more severely in HOM mice of both sexes, however, male mice had more similarly associated phosphopeptides that increased than females, suggesting increased basal signaling of ΔF48 PKCγ. For both sexes, ontologies were centered on neuronal and cytoskeletal processes. This could reflect changes in Purkinje cell morphology in SCA14 genotypes. Although ontologies were conserved, the signaling mechanisms involved may be different as phosphopeptides were not highly conserved between comparisons.

To further explore sex differences in the phosphoproteome, we compared SCA14 genotypes between male and female mice. In HET mice, volcano plots identified phosphopeptides that were significantly increased (more abundant in male HET) or decreased (more abundant in female HET) (**Figure 6a**). Corresponding gene ontology (GO) analysis was performed for significantly different phosphopeptides found in HET male compared to HET female mice for biological processes (**Figure 6b**), cellular components (**Supplemental Figure 5m**), and molecular functions (**Supplemental Figure 5n**). The top 10 most significantly enriched for ontologies for biological processes are shown (p<0.05). Gene ontology analysis was also performed for phosphopeptides that were associated in both sexes. Sex similarly associated phosphopeptides were designated as phosphopeptides that had the same directionality of change in HET vs WT in both sexes and were significantly different in at least one sex (biological processes, **Figure 6c**; cellular components, **Supplemental Figure 5o**; molecular functions, **Supplemental Figure 5p**). These sex similarly associated phosphopeptides are displayed as heatmaps, with increasing phosphopeptides (**Figure 6d**) being more prevalent than decreasing phosphopeptides (**Figure 6e**). As with the HET mice, volcano plots (**Figure 6f**), gene ontologies (**Figure 6g-h; Supplemental Figure 5q-t**), and heatmaps (**Figure 6i-j**) are displayed for HOM male mice compared to HOM females. As in HET mice, sex similarly associated phosphopeptides are displayed as heatmaps with increasing phosphopeptides (**Figure 6i**) being more prevalent than decreasing phosphopeptides (**Figure 6j**). Of the differentially abundant phosphopeptides in HET mice, 41% are similarly altered in both sexes, and 43% are similarly altered in both sexes in HOM mice, suggesting that the majority of altered phosphopeptides in each genotype are not commonly altered in both sexes. Ontologies for these comparisons were centered on neuronal and cytoskeletal processes for both different and similarly altered phosphopeptides. Notably, the most significant sex-specific changes were associated with synaptic functions. Although overlap in ontologies occurred between comparisons, specifically altered phosphopeptides remained unique between all groups, suggesting that similar biological processes were affected, but not necessarily through the same pathway.

**Figure 6:**
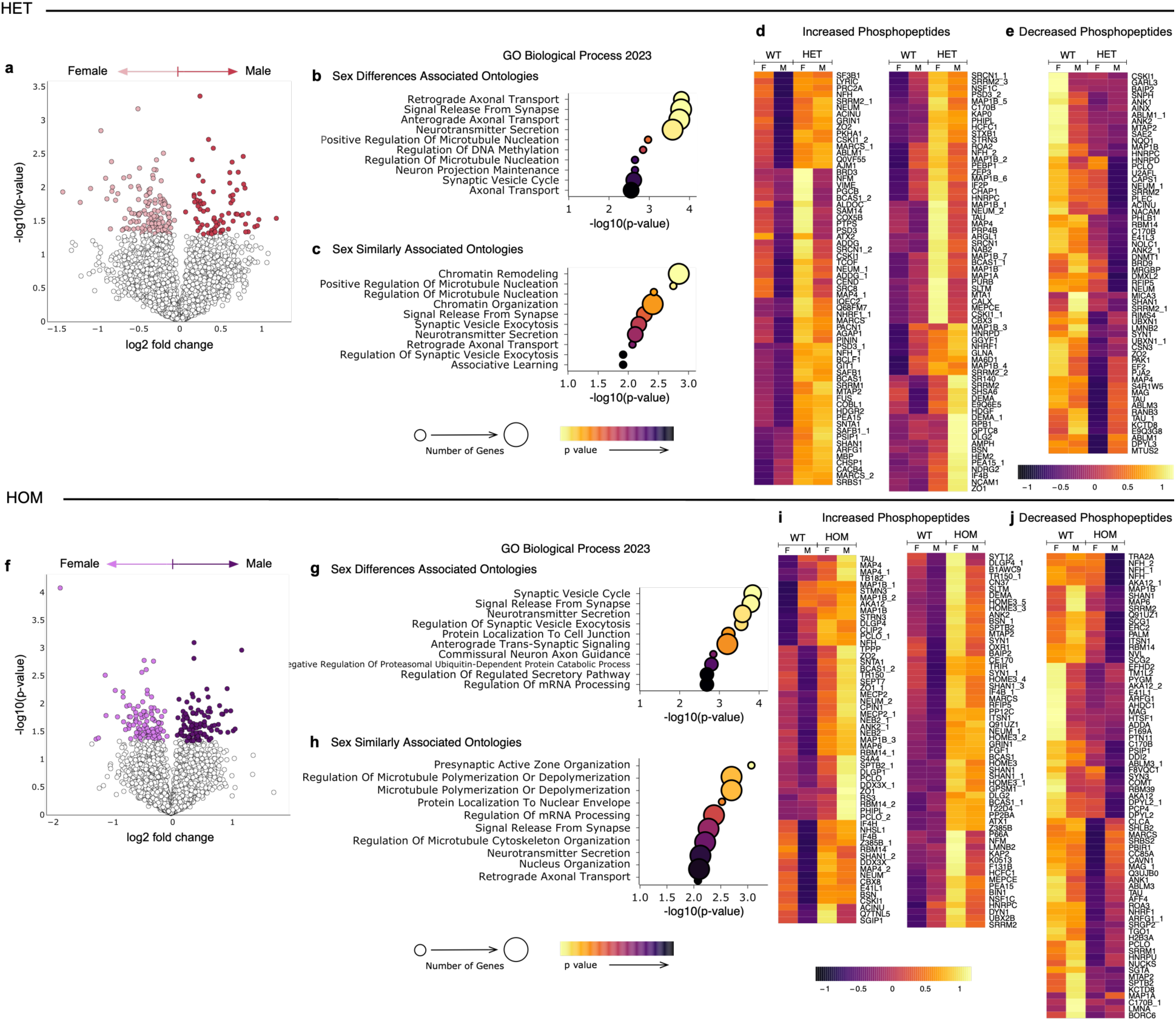
Sex differences in the SCA14 mouse cerebellar phosphoproteome. Phosphoproteomic analysis was carried out as in Figure 5. **a)** Volcano plots show change in phosphopeptide abundance between male HET and female HET mice. Color represents phosphopeptides with p-value < 0.05. Dot plots indicate the top 10 most significantly enriched for gene ontologies from the differentially abundant phosphopeptides identified in **b)** male HET vs female HET mice, and **c)** similarly altered phosphopeptides in HET mice (defined as phosphopeptides with the same trend in males and females with a significant difference in intensity between WT vs HET in at least one sex). Similarly altered phosphopeptides are shown in heatmaps separated by those that **d)** increase or **e)** decrease in HET mice compared to WT. **f)** Volcano plots show change in phosphopeptide abundance between male HOM and female HOM mice. Color represents phosphopeptides with p-value < 0.05. Dot plots indicate the top 10 most significantly enriched for gene ontologies from the differentially abundant phosphopeptides identified in **g)** male HOM vs female HOM mice, and **h)** similarly altered phosphopeptides in HOM mice (defined as phosphopeptides with the same trend in males and females with a significant difference in intensity between WT vs HOM in at least one sex). Similarly altered phosphopeptides are shown in heatmaps separated by those that **i)** increase or **j)** decrease in HOM mice compared to WT. Dot plot color scale indicates p-value, dot size indicates number of genes per ontology (p<0.05). Heatmap color scale indicates normalized phosphopeptides intensity with values centered and scaled by row. Protein names accompanied by a number designate multiple phosphorylation sites for the same protein.

## Discussion

In this study, we have developed a new SCA14 mouse model which contains the amino acid deletion ΔF48 of PKCγ identified in human patients. This mutation was selected because of the severity of the biochemical defect and resulting severity of the associated symptomology(11, 32). As with other SCA14 mutations, ΔF48 PKCγ has impaired autoinhibition that results in ‘leaky’ basal activity but is unique in that this mutation breaks communication between the DG sensor (C1B domain) and the autoinhibitory pseudosubstrate. As a result, it is unresponsive to agonist-evoked activation(21, 32, 33), the downstream effects of which are unknown. Thus, the mutation provides a unique tool to examine how leaky basal activity, associated with SCA14 mutations, drives the disease. Behavioral, immunohistochemical, phosphoproteomic, and biochemical analysis revealed motor dysfunction, altered Purkinje cell morphology, a rewired phosphoproteome with notable effects in neuronal and cytoskeletal processes, and no overall change in PKC substrate phosphorylation despite significantly reduced levels of PKCγ. Strikingly, phenotypes were significantly more severe in males suggesting a potential neuroprotective mechanism in female SCA14 mice that may also be reflected in SCA14 human patients. Our data reveal that the aberrant signaling of ΔF48 PKCγ is sufficient to drive ataxia related phenotypes, with male mice displaying greater biochemical, cellular, and behavioral aberrations.

### SCA14 mice display sex differences that indicate a potential neuroprotective mechanism in female mice

A major finding in this study was the evidence of sex differences in our SCA14 mouse model with ΔF48 PKCγ. Little information is available on sex differences in SCA14 human patients or models. This finding highlights an important factor to consider when developing treatment strategies in SCA14 as well as other neurodegenerative diseases associated with PKC activity such as Alzheimer’s disease(65, 66). We found that SCA14 male mice displayed more severe differences compared to WT in most of our inquiries including ataxia-related behaviors, cerebellar morphology deficits, aberrant PKCγ expression, and alterations in the phosphoproteome. Indeed, sex-specific differences were as significant as genotype-specific differences vis a vis alterations in the phosphoproteome. Additionally, for some tests, HET male mice displayed differences that were not observed in the HET female mice. We also noted reduced viability of HOM male mouse offspring, detected as a lack of HOM male mice born. Thus, at least in this SCA14 mouse model, male mice are much more impacted than female mice.

Having observed strong sex differences in our SCA14 mice, we investigated human patient data for differences in disease severity in male and female patients with SCA14. Using the Online Mendelian Inheritance in Man (OMIM, #605361) database, we analyzed the available data of 77 patients with SCA14 that had clinical descriptions that included the age of onset and sex of the individuals(5, 13, 67-70). Since ΔF48 PKCγ is a rare mutation that is lacking in patient data, we used data from individuals with a variety of PKCγ mutations causative for SCA14. Interestingly, the reported age of onset of SCA14 symptoms in female patients was significantly higher (37 years old) than in male patients (29 years old) (**Figure 7a-b**). Additionally, fewer male patients were identified than female, which could reflect a loss in viability of male humans (30 males and 47 females; **Figure 7a**). This is consistent with our findings from the mouse model suggesting that female patients may have neuroprotective mechanisms that diminish the start of symptomology and increase viability. Thus, despite a relatively small sample size in the human cohorts, SCA14 appears to affect males more profoundly than females across various SCA14 mutations. Future attention to sex differences may provide more insight into how general sex differences are in the different SCA diseases.

**Figure 7:**
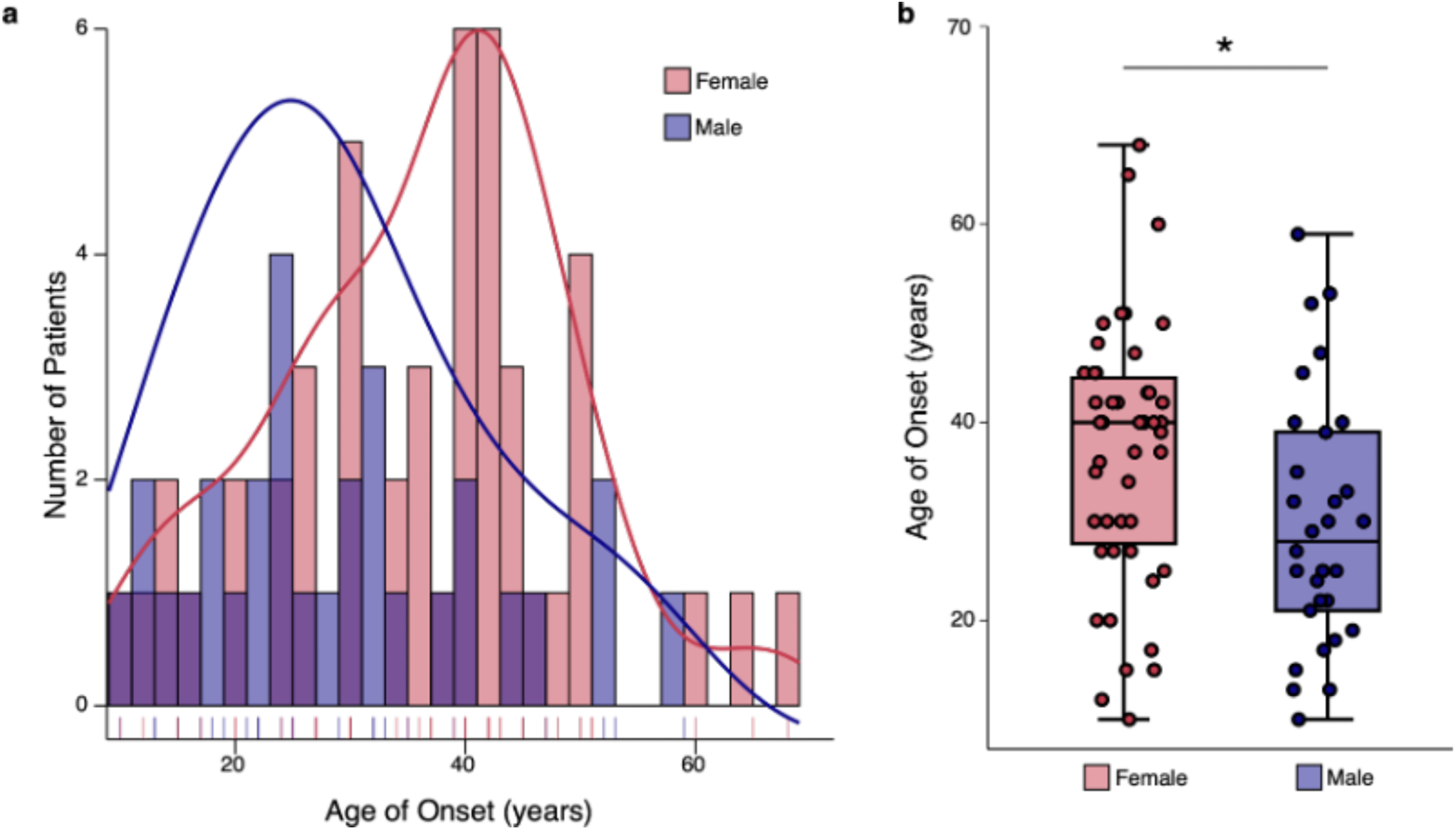
SCA14 has a later age of onset in female patients compared to male patients. Using the Online Mendelian Inheritance in Man (OMIM) database, data from 77 SCA14 patients was obtained from studies that reported a clinical description that included the age of onset and sex. **a)** Histogram displays the age of onset reported by SCA14 patients with sex designated by color, and the density curve shows the distribution for female patients (red) and male patients (blue). **b)** Box plot displays the significant difference in age of onset between female patients (red) and male patients (blue) as determined by two sample t-test (p= *<0.05). Mean±S.E.M.: Female=37±1.9, Male=29±2.3.

Given the decreased severity of SCA14 symptoms in females, an intriguing possibility is that estrogen, particularly 17-β estradiol (E_2_), may provide a neuroprotective effect. Consistent with this, E_2_ has been shown to be neuroprotective in several diseases, including cerebellar ataxia models(71-79). Furthermore, inhibition of PKC has been reported to prevent E_2_ neuroprotection of neuronal cultures(78). E_2_ is dominantly expressed in females during reproductive years and is involved in a wide range of physiological and behavioral functions. E_2_ is an agonist for transcription-factor estrogen receptors (ERα/ERβ) and G-protein coupled estrogen receptors (GPER1), modulating gene regulation and intracellular signaling respectively(45, 79-81). Notably, estrogen has been reported to increase PKC activity and/or expression in a variety of tissue and cell types including brain(82-89). E_2_ induces activation of PLC, thought to be through GPER1 signaling. Activation of PLC increases the production of DG/Ca^2+^ and thus agonist-evoked activation of PKCγ(87-90). ERs are abundant in cerebellum, and ERβ and GPER1 are dominantly expressed in Purkinje cells(45, 91-95). E_2_ has also been shown to modulate Purkinje cell function(45, 96, 97) and is biosynthesized in the brain(43, 71, 77, 79, 98). This evidence, combined with our data, implicates E_2_ as a potential neuroprotective agent for SCA14. Since ΔF48 PKCγ is both gain-of-function (increased basal activity) and loss-of-function (impaired agonist-evoked signaling and loss of dynamics of activity), perhaps E_2_ can increase cellular DG/Ca^2+^ to enhance signaling by any functional PKC (i.e. WT PKCγ/α) in the cerebellum. This may overcome aberrant ΔF48 PKCγ signaling and allow WT PKCγ to mask the deleterious effects of ΔF48 PKCγ in females. Future research on the role of E_2_ on PKC signaling in SCA14 is warranted.

### Increased basal activity of ΔF48 PKCγ compensates for loss of total PKCγ

Our investigation of PKCγ expression and localization revealed a dramatic gene dose-dependent loss of total PKCγ in both males and female mice. We have previously shown that the cellular turn-over of ΔF48 PKCγ is an order of magnitude faster than that of WT PKCγ because of inherent instability of the mutant PKCγ(21, 32). Thus, it is not surprising that steady-state levels of ΔF48 PKCγ are low in HET and HOM mice. Aberrant PKC dimerizes with WT enzyme, trapping it in a degradation-sensitive conformation; therefore, levels of WT PKCγ and its paralog PKCα may also be reduced. Unlike PKCγ, PKCα is expressed in other cell types in the cerebellum which may explain why results from the Western blot and immunofluorescence analysis for PKCα differed. Importantly, the unrestrained activity of the ΔF48 PKCγ is sufficient to drive robust substrate phosphorylation. A similar finding was reported for another SCA14 mouse model: a transgenic mouse expressing a pseudosubstrate mutant PKCγ, A24E, displays a 10-fold reduction in PKCγ steady-state levels yet results in a paradoxical increase in cerebellar substrate phosphorylation(34). It is noteworthy that deletion of the PKCγ gene does not produce some SCA phenotypes such as synaptic transmission and neurodegeneration(99). This reveals that it is not loss of PKCγ, but the presence of aberrant PKCγ that drives some ataxia-associated phenotypes. These findings underscore the importance of unrestrained ‘leaky’ activity of PKCγ in driving SCA14.

Little is known about PKCγ expression in SCA14 human patients, however, one study of a male patient with SCA14 reported reduced PKCγ levels in Purkinje cells. This affected person was a 66-year-old male who died of an aneurysm with the SCA14-associated mutation H101Y in PKCγ. Immunohistochemistry of cerebellar tissue revealed a loss of Purkinje cells; however, the remaining Purkinje cells (which had low PKCγ staining) had equivalent Calbindin D28k staining as control cerebellum, suggesting the loss of PKCγ in Purkinje cells was unrelated to Purkinje cell death in this SCA14 patient(2). Kapfhammer and colleagues have previously suggested that it is not Purkinje cell loss, but rather Purkinje cell dysfunction, that drives ataxic phenotypes(53).

### ΔF48 PKCγ causes cerebellar Purkinje cell dysfunction rather than loss in SCA14 mice

SCA14 pathogenesis can be driven by neurodegeneration of Purkinje cells, or by dysfunction and altered physiological activity of Purkinje cells that disrupt cerebellar signaling leading to motor dysfunction(6, 7, 34, 48, 56, 57, 58). Although neurodegeneration is a hallmark of SCA, Purkinje cell dysfunction and deficits in morphology have been observed in the absence of Purkinje cell death in SCA14 models, and in some cases Purkinje cell dysfunction precedes ataxic phenotypes(34, 48, 49, 53, 56). Additionally, some SCA14 cases in human patients only show mild cerebellar atrophy, suggesting neurodegeneration may not be the determinant for SCA14 progression(12, 53). Our mouse model of SCA14 is consistent with this concept. We did not observe overt Purkinje cell loss, rather a more subtle alteration in Purkinje cell morphology that is indicative of Purkinje cell dysfunction. Additionally, histology and behavioral tests were performed at multiple ages (5-10 months) and combined because no differences were observed in neurodegeneration of ataxic behaviors with increasing age. Future studies will consider younger animals to assess age differences through development. These morphological changes may be due to developmental deficits in Purkinje cell arborization, similar to previous studies that showed that enhanced PKC activity compromises dendritic growth(48-53). How aberrant PKC signaling causes neurodegeneration and deficits in Purkinje cells remains to be established, therefore, additional research is needed to further investigate the role of PKC in Purkinje cell development as well as mature Purkinje cell function and signaling in the context of SCA14. Importantly, these results present other avenues of interest in developing therapies for specific SCA14 patients, where restoring Purkinje cell function may be more efficacious than counteracting neurodegeneration.

### Phosphoproteomic analysis indicates altered neuronal and cytoskeletal processes in SCA14 mice unique to genotype and sex

Phosphoproteomic analysis of the cerebellum of ΔF48 PKCγ mice identified significantly rewired signaling in both male and female mice, with many affected processes related to neuronal signaling. Notably, differentially abundant phosphopeptides identified in SCA14 genotypes were significantly enriched for GO Biological Processes including Neurotransmitter Secretion, Signal Release From Synapse, and Retrograde and Anterograde Axonal Transport. Likewise, enriched for GO Cellular Components included neuronal associated components such as Neuron Projection, Synaptic Membrane, and GABA-ergic Synapse. Alterations in phosphopeptides associated with neuronal processes likely reflect disrupted signaling associated with the observed alterations in Purkinje cell neurons in SCA14 mice. This further implicates Purkinje cell and cerebellum dysfunction in our SCA14 mice despite a lack of Purkinje cell death.

In addition to neuronal related processes, cytoskeletal-associated ontologies were enriched in the differentially abundant phosphopeptides identified in SCA14 genotypes, for example significantly enriched for GO Biological Processes included Regulation Of Microtubule Polymerization Or Depolymerization, Cytoskeleton Organization, and Membrane Organization in SCA14 genotypes. Likewise, significantly enriched for GO Molecular Functions for cytoskeletal associated ontologies included Tubulin Binding, Microtubule Binding, and Actin Binding. PKC is a known regulator of the cytoskeleton and microtubule dynamics, therefore dysregulation of PKCγ will likely disrupt signaling in cytoskeletal associated pathways as observed in our SCA14 mice(22). The mice with the SCA14-associated mutation H101Y PKCγ mutation also have alterations in phosphorylation of cytoskeletal related proteins identified via phosphoproteomic analysis, including neurofilament proteins that play key roles in growth of axons, with aberrations associated with neurodegeneration(32, 100). We have observed similar alterations in the phosphorylation of cytoskeletal associated proteins including neurofilament proteins (NFH, NFL, NFM) in our SCA14 mice, suggesting that ΔF48 PKCγ dysregulates similar signaling pathways involved in the regulation of neuron cytoskeletal structures and functions. Moreover, cytoskeleton dynamics in the brain are critically involved in neuron and synapse formation and maintenance(22), further implicating dysregulated PKCγ signaling and phosphoproteomic rewiring as an underlying cause of Purkinje cell and cerebellum dysfunction.

Although ontologies centered on neuronal and cytoskeletal processes, specific ontologies were not necessarily enriched for in all comparisons between genotype and sex. Additionally, specific ontologies that were enriched for in multiple comparisons had a limited number of shared differentially abundant phosphopeptides between groups. For example, the GO biological process Regulation Of Microtubule Polymerization Or Depolymerization was found to be significantly enriched for in all SCA14 genotypes compared to WT; however, not one phosphorylation site found on proteins in this ontology was the same in all comparison including both sexes. Therefore, although the same ontologies and pathways are altered in SCA14 mice, the underlying mechanism may differ based on sex and genotype, adding further complexity to SCA14.

A major finding in our data was that the aberrant signaling of ΔF48 PKCγ was sufficient to rewires the cerebellar phosphoproteome uniquely by genotype and sex. Notably, male HOM mice showed the greatest number of significantly altered phosphopeptides, supporting our hypothesis that males with SCA14 are affected more severely. Importantly, of the identified HET and HOM similarly associated phosphopeptides in SCA14 male mice, the majority were increased (64%), and male mice had 67 more phosphopeptides increased compared to females. This may suggest a higher degree of overactivity of ΔF48 PKCγ in male mice and more deficits in signaling pathways associated with PKCγ. This was also reflected in the sex similarly associated phosphopeptides, where in both HET and HOM mice, more phosphopeptides were increased than decreased, suggesting a higher degree of variability in phosphopeptides that were reduced in SCA14 genotypes between sexes. Phosphopeptides that were significantly decreased in SCA14 genotypes compared to WT, despite the overactive ΔF48 PKCγ, likely reflects PKC regulation of phosphatase function to reduce phosphorylation of some proteins(101).

When comparing HET and HOM similarly altered phosphopeptides, the majority of phosphopeptides trended the same within sex, suggesting that HOM mice may have similar, but more severe alterations in the phosphoproteome as HET mice. These findings add validity and biological relevance to our SCA14 mouse model, where HOM animals may be used in studies for a more robust and clear readout while still representing the HET mice, as human SCA14 is autosomal dominant. This is also reflected in our behavioral and molecular studies, where HOM mice often displayed a robust, significant alteration while HET mice displayed a trend in the same direction, with the exception that female mice often had confounding results potentially due to neuroprotective mechanisms. Therefore, our SCA14 mouse may be a good representation for continued experimentation including future research to modulate PKCγ signaling and SCA14 phenotypes. Furthermore, the limited number of individuals identified with ΔF48 PKCγ(11) occludes establishment of typical disease progression or sex differences, warranting a model to study its underlying mechanism. This model may also shed light on other SCA14 mutations, or other neurodegenerative diseases, that present dysregulated PKCγ signaling(65, 66).

In summary, our study identifies sex-specific differences in the biochemical, cellular, and behavioral effects of the ΔF48 PKCγ SCA14 mutation, with males significantly more affected than females. Further studies are needed to decipher the molecular mechanisms driving these differences, with one possible mechanism being protective effects of estrogen. Our data suggest that treatment strategies to reduce aberrant ΔF48 PKCγ may be particularly beneficial in males.

## Methods

### Sex as a biological variable

This study examined male and female mice in all genotypes (WT, HET, HOM). Sex-dimorphic effects are reported for the SCA14 mouse model.

### Generation of transgenic mice

#### CRISPR target sequence design

gRNAs were designed using available online tools such as CRISPR RGEN Tool (http://www.rgenome.net/) and CRISPRDirect (https://crispr.dbcls.jp). gRNAs were prepared via the annealing of crRNA (5’ GTCGGTACAGTGACTGCAGA 3’) and tracrRNA (Alt-R® CRISPR-Cas9 tracrRNA; cat. No. 1072532) from Integrated DNA Technologies (Coralville, IA, USA). The crRNA and tracrRNA were chemically synthesized, and RNase-Free HPLC purified by Integrated DNA Technologies (Coralville, IA, USA). Single-strand oligonucleotide (ssODN, 5’ AGAGCCACAAGTTCACCGCTCGTTTCTTCAAGCAGCCAA**CAT**GCAGTCACTGTACCGACTT CATCTGGTGAGGGAAGCGGGCTAGGGGAG 3’) was chemically synthesized and standard desalted by GENEWIZ (South Plainfield, NJ, USA).

#### Cas9 protein

Cas9 protein (Alt-R S.p. Cas9 Nuclease V3; cat. no. 1081058) was obtained from Integrated DNA Technologies (Coralville, IA, USA)

#### Mixiture preparation for the microinjection

Cas9 protein, the duplex of crRNA and tracrRNA, and ssODN were diluted and mixed in IDTE buffer from Integrated DNA Technologies (Coralville, IA, USA) to a working concentration of 30 ng/μl, 0.6pmol/μl, and 10 ng/μl, respectively. The mixture was incubated at room temperature for 10 minutes.

#### Embryo recovery, microinjection, and embryo implantation

Pronuclear-stage embryos were recovered from the superovulated 3-5 weeks old C57BL/N females by the intra-peritoneal (IP) injection of pregnant mare serum gonadotropin (PMSG) at 5 units/mouse (cat. no. HOR-272, Prospec, East Brunswick, NJ, USA) followed by human chorionic gonadotropin (hCG, 5 units/mouse, cat. no. C8554, Millipore Sigma, St. Louis, MO, USA) over 46-48 hours interval later and then mated overnight with C57BL/N males. Pronuclear-stage embryos were harvested 16 hours after HCG injection. The ampulla region of oviducts was washed in M2 medium (EmbryoMax M2 medium; cat. no. MR-015-D, Millipore Sigma, St. Louis, MO, USA) and transferred into 0.5 mg/ml of hyaluronidase (cat. no. H4272, Millipore Sigma, St. Louis, MO, USA). The eggs were released by the punch with needle. After 1 minute, the cumulus cells were separated, and the pronuclear-stage embryos were washed in fresh M2 medium three times and incubated in 30μl of M2 medium at 37°C until the microinjection is completed. For microinjection, the pronuclear-stage embryos were transferred to 100 μl of M2 medium in a cover glass and microinjected with 1∼2 picoliters of the mixture using an inverted microscope (Eclipse Ti, Nikon, Tokyo, Japan) equipped with a Femtojet express microinjector (Eppendorf, Enfield, CT, USA). Microinjected pronuclear-stage embryos were transferred into the ampulla of pseudo-pregnant females.

### Genotyping

Tissue obtained from mouse ear clips was digested with tail lysis buffer (1 M Tris pH8, 5 M NaCl, 0.5 M EDTA, 10% SDS), and genomic DNA was isolated and amplified by PCR (forward primer 5’-AGGTGCTGAGAGCGAAGC-3’ and reverse primer 5’-GGGTGAAATTGGGAAACTGC-3’), using Q5 PCR Master Mix (New England Biolabs, cat. no. M0492). DNA was subjected to restriction enzyme digestion by NspI (NspI restriction digest site is present only in ΔF48 PKCγ alleles) in CutSmart Buffer (New England Biolabs, cat. no. B6004) and visualized via electrophoresis on a 0.8% agarose gel to determine the genotype [WT/WT (WT), WT/ΔF48 (HET), or ΔF48/ΔF48 (HOM)].

### Behavior testing

For behavioral experiments: Mice were acclimated to the procedure room prior to testing. Mice were returned to their home cage at the end of the test session. The arena/equipment was wiped down with 70% ethanol before the start of each run. Experimenter was blind to genotype for all tests performed, and each group (Male WT, Male HET, Male HOM, Female WT, Female HET, Female HOM) contained animals from at least three separate litters. Each test was performed at least twice with a minimum of 24 hours rest between each test session, and data are averaged across trials. Multiple cohorts were tested, and all groups were represented in each cohort. No differences were observed based on the age of the mice tested, and ages were combined (age=5-10 months). Sample sizes reflect availability of mice per genotype (N=5-16). Data are normalized within cohort to the respective WT group when combined to avoid batch effects.

#### Rotarod test

To test locomotor coordination and balance, the Rotarod test was performed by placing mice on 3cm rotating drum (Accurotar, Omnitech Electronics, Inc., Columbus, OH) and measuring the latency to fall. The rod started in a stationary state and then began to rotate with a constant acceleration of 10 rpm. Three trials were performed for four days and averaged per animal.

#### Treadmill walking test

For this test, mice were first trained to walk on the treadmill (Columbus Instruments Exer-3/6, Columbus, OH) in three daily 5 min sessions in which stopping resulted in the mice touching the back of the apparatus and experiencing a mild shock (200 msec pulses of electric current with pulse repetition rate of 3 times per second (3 Hz) and an intensity of 1 mA). One week later, the mice were run at a speed of 10 cm/s for one minute without receiving shock and the percentage of time the mice walked ahead of the bumper/rear of the apparatus was assessed. Behaviors such as sliding, facing backwards, or climbing were subtracted.

#### Ladder rung test

To assess skilled walking and coordination, mice were first trained to walk across a horizontal ladder with evenly spaced rungs (1cm apart) to reach their home cage, then tested three additional times with the spacing of the rungs randomized (i.e. rungs removed at random, with more rungs removed in each subsequent trial). The number of times the mouse’s paw slipped from a ladder rung was counted (from video recordings) and averaged over the three random trials.

#### Wire hang test

The hanging wire test allows for the assessment of grip strength and motor coordination. A 2 mm wire was securely suspended between two stands at a height of 37 cm. Padding was placed in a cage under the wire to safely catch the mice after they fall. Mice were suspended by the tail and gently lowered until they grasped the wire with their front paws. The tail was then released, and the mice were allowed to hang from and move on the wire freely. The mice were timed until they fell from the wire (maximum 120 seconds per trial). The latency to fall time was measured and averaged over two trials.

#### Grip strength test

Forelimb grip strength was acquired with a digital Grip Strength Meter (Columbus Instruments Model 0167-004L, Columbus, OH). Each mouse was held so that only the forelimb paws grasped the flat mesh assembly and then pulled back until its grip was broken. The peak force reached was averaged across three consecutive trials.

### Real-time qPCR

Cerebellum samples from WT, HET, and HOM male and female mice at 6 and 8 months old were dissected, flash frozen in liquid nitrogen, and cryo-pulverized on dry ice using a stainless-steel mortar and pestle. Crushed cerebellum was transported to a 1.5 ml tube. RNA was extracted using TRI reagent (Sigma-Aldrich, cat. no. T9424). The TRI reagent protocol was followed for RNA isolation or extraction per manufacturers recommendation and resuspended in RNase free water (TRI reagent; Sigma-Aldrich, cat. no. T9424). RNA was assessed with a Nanodrop spectrophotometer (Thermo Fisher Scientific). Next, cDNA was synthesized using SuperScript III first-strand synthesis system for RT-qPCR with 500 ng of RNA input (Invitrogen, cat. no. 108080-051). A control sample lacking reverse transcriptase was included for each sample to assess gDNA contamination. RT-qPCR was run using TB Green premix Ex Taq II Kit (TAKARA, cat. no. RR820A) on QuantStudios III PCR (Applied Biosystems). Standard curves were used to assess efficiency and detection limits of primer sets and determine the appropriate quantity of RNA before testing samples (PKCγ: forward primer 5’-ACCAGGGCATCATCTACAGG-3’ and reverse primer 5’-CTTCCTCATCTTCCCCATCA-3’; GAPDH: forward primer 5’-AGGTCGGTGTGAACGGATTTG-3’ and reverse primer 5’-TGTAGACCATGTAGTTGAGGTCA-3’). Samples were run in triplicate and the ΔΔCt method was used to analyze the data. No significant differences were observed between male or female WT samples. Experimenter was blind to genotype and sex where applicable. Graphs represent mean±S.E.M. normalized to the respective WT control group, N=5-9 mice.

### Immunofluorescence

#### Tissue preparation

Mice were anesthetized (isoflurane) and perfused with cold PBS followed by freshly prepared 4% paraformaldehyde (PFA) (age=5-10 months). Mice were decapitated and skulls were immersion fixed in 4% PFA overnight at 4°C, followed by brain dissection and an additional immersion fixation in 4% PFA at 4°C for 1 hour. Brains were then cryoprotected by immersing them in 30% sucrose (w/v) in PBS at 4°C until they sink. Brains were then placed in cryomolds (Polysciences Inc., cat. no. 18646D-1) with OCT (VWR, cat. no. 25608-930). Molds were placed into a dry ice/ethanol bath until frozen. Brains were demolded and stored at -80°C until sectioning by the UCSD histology core facility. Brains were sectioned sagittaly into 40 μm sections using a cryostat and kept at -20°C floating in cryoprotectant media.

#### Immunostaining

The sections were treated in a 24-well plate on a shaker. They were first treated with 0.3% Triton X-100 in PBS for 10 minutes to remove lipids, then treated with a blocking solution (3% donkey serum, 1% BSA, 1% Fish Gelatin, 0.1% Triton X-100, 50 mM Glycine in PBS) for 1 hour in room temperature. Afterward, they were incubated with antibodies as listed below, sequentially, in cold room overnight. The first antibody is rabbit anti-PKCγ (GeneTex, cat. no. GTX107639), the second antibody is mouse anti-Calbindin D28k (Swant D28k, cat. no. CB300), the third antibodies are a combination of AF568 Donkey anti-Rabbit IgG (Invitrogen, A10042) plus AF680 Donkey anti-mouse IgG (Invitrogen, cat. no. A10038) plus Cy2 Donkey anti-Guinea pig IgG (Jackson immune research, cat. no. 706-225-148), and the last antibody is AF790 Rabbit anti-PKC alpha (Santa Cruz Bio, cat. no. sc-8393). Finally, the samples were incubated with freshly prepared 100 ng/ml DAPI in PBS for 5 minutes before mounted on glass with a water-based medium (SouthernBiotech, cat. no. 0100-20). Confocal images were obtained with a Nikon eclipse Ti2 microscope in UCSD Nikon Image Center with a 20x lens with a numerical aperture of 0.75. Z-stacks were taken through the whole cerebellar section at 2 µm intervals, and the entire cerebellum was tile scanned and stitched together. Microscope settings were kept the same for all slices during image acquisition.

#### Image analysis

Image analysis was performed using FIJI (ImageJ version 2.14.0/1.5f)(102) to quantify the PCL and ML of images. The cerebellar image z-stacks were average projected. Next, a Region of Interest (ROI) was created for the PCL and ML, using a mask for all areas expressing Calbindin D28k, created by auto-thresholding Calbindin D28k images with manual adjustment to the ROI if needed to only select for the PCL and ML. Mean fluorescent intensity was measured within this ROI to obtain relative levels of immunofluorescent staining intensity for each protein interrogated. Background was also measured and subtracted per sample. Mean fluorescent intensity was normalized to WT controls within imaging sessions. The ROI was used to unbiasedly obtain the thickness of the PCL and ML by using the Local Thickness plugin(103). To obtain the linear density of Purkinje cells, the PCL was then traced through all lobules, and Analyze Particle was used to unbiasedly and automatically count the Purkinje somas along the traced selection. Purkinje cell counts were then divided by the selection distance to normalize for variations in distance. Experimenter was blind to genotype and sex for all steps in the immunostaining process when possible. Data is represented as mean±S.E.M. and intensity values were normalized to the respective WT control group, N=3-6 mice. No significant differences were observed between male or female WT samples. All image analysis software settings were kept the same between samples and only raw images were analyzed. Brightness and contrast were adjusted in the representative images and all represented images were adjusted the same.

### Western blot analysis

Cerebellum samples from WT, HET, and HOM male and female mice at 6 and 8 months old were dissected, flash frozen in liquid nitrogen, and cryo-pulverized on dry ice using a stainless-steel mortar and pestle. Crushed cerebellum was transported to a 1.5 ml tube and lysed in Tris-lysis buffer (1 M Tris pH 7.4, 10 mM Sodium Pyrophosphate, 50 mM sodium fluoride, 5 mM EDTA, 1% Triton X-100) and briefly sonicated. Lysis buffer was supplemented with 1 mM PMSF, 50 µg/ml leupeptin, 1 mM Na_3_VO_4_, 2 mM benzamidine, 1 µM microcystin, and 1 mM DTT added immediately prior to lysis. Protein concentration was quantified by BCA (cat. no. 23227, Pierce). Samples were boiled in sample buffer (250 mM Tris HCl, 8% (w/v) SDS, 40% (v/v) glycerol, 80 µg/ml bromophenol blue, and 2.86 M β-mercaptoethanol) for 5 minutes at 95°C. SDS-PAGE was performed on protein samples (10-20 µg) using 10% acrylamide gels (bis/acrylamide solution, cat. no. 161-0156, BioRad) with protein ladder (cat. no. 161-0394, BioRad). Gels were transferred to 0.45 µm nitrocellulose membranes (cat. no. 1620115, BioRad) by a wet transfer method at 4°C for 1 hour at 100 V in transfer buffer (200 mM Glycine, 25 mM Tris Base, 20% Methanol). Membranes were blocked in 5% (w/v) nonfat dry milk in phosphate buffer saline with tween-20 (PBS-T) for 1 hour at room temperature, then incubated in primary antibodies diluted in 1% BSA in PBS-T overnight at 4°C. Primary antibodies were against the following: PKCα (1/1000), pSer PKC substrate (1/1000), PKCγ (1/1000), α-Tubulin (1/10,000), Calbindin D28k (1/1,000), PKCδ (1/1000), Vinculin (1/1000), phospho-GSK3 α/β(Ser21/Ser9) (1/1000), total GSK3 α/β (1/1000), phospho-MARCKS (Ser159/Ser163) (1/1000), total MARCKS (1/10,000), PKCη (1/1000), or GAPDH (1/5000). Membranes were washed for 5 minutes three times in PBS-T and incubated with goat anti-rabbit 800 nm and or goat anti-mouse 700 nm secondary antibodies (1/10,000 in 1% BSA in PBS-T) for 1 hour at room temperature, then imaged using Azure Biosystems Sapphire FL Biomolecular Imager and raw images were quantified with FIJI (ImageJ version 2.14.0/1.5f). No significant differences were observed between male or female WT samples. Experimenter was blind to genotype and sex where applicable. Graphs represent mean±S.E.M. normalized to the respective WT control group within gel, N=6-11 mice.

### Antibodies

Antibodies used are listed with the company from which they were purchased and catalog number: PKCα (BDT, cat. no. 610108), pSer PKC substrate (Cell Signaling, cat. no. 2261, lot 26), PKCγ (Santa Cruz, cat. no. C-19; or GTX, cat. no. 107639), PKCα (BD Transduction, cat. no. 610108; or Santa Cruz Bio, cat. no. sc-8393), α-Tubulin (Sigma Aldrich, cat. no. T6074), Calbindin D28k (Swant, cat. no. 300), PKCδ (BD, cat. no. 610397), phospho-GSK3 α/β(Ser21/Ser9) (Cell Signaling, cat. no. 9331), total GSK3 α/β (Cell Signaling, cat. no. 9832), phospho-MARCKS (Ser159/Ser163) (Cell Signaling, cat. no. 11992), total MARCKS (Proteintech cat. no. 20661), Vinculin (Cell Signaling, cat. no. 4650), PKCη (Abcam, cat. no. 179542), GAPDH (Cell Signaling, cat. no. 2118), Goat anti-mouse, Azure700 conjugate (Azure Biosystems, cat. no. AC2129), and Goat anti-rabbit, Azure800 conjugate (Azure Biosystems, cat. no. AC2134).

### Phosphoproteomics

#### Sample preparations

The cerebellum of WT, HET, and HOM male and female mice at 6 months old was isolated, flash frozen in liquid nitrogen, and cryo-pulverized. Tissue was sent to the UCSD Biomolecular and Proteomics Mass Spectrometry Facility for Phosphoproteomic analysis (N=3 mice). Experimenter was blind to genotype and sex of samples. Samples were lyophilized overnight and reconstituted in 200 µl of 6 M Guanidine-HCl. The samples were then boiled for 10 minutes followed by 5 minutes cooling at room temperature. The boiling and cooling cycle was repeated a total of 3 cycles. The proteins were precipitated with addition of methanol to final volume of 90% followed by vortex and centrifugation at maximum speed on a benchtop microfuge (14000 rpm) for 10 minutes. The soluble fraction was removed by flipping the tube onto an absorbent surface and tapping to remove any liquid. The pellet was suspended in 200 µl of 8 M Urea made in 100 mM Tris pH 8.0. TCEP was added to final concentration of 10 mM and Chloro-acetamide solution was added to final concentration of 40 mM and vortex for 5 minutes. 3 volumes of 50 mM Tris pH 8.0 were added to the sample to reduce the final urea concentration to 2 M. Trypsin (1:50 ratio) was incubated at 37°C for 12 hours. The solution was then acidified using TFA (0.5% TFA final concentration) and mixed. The sample was desalted using C18-StageTips (Thermo Fisher Scientific) as described by the manufacturer protocol. The peptide concentration of sample was measured using BCA. 80ug of each sample were used in TMT labeling. Thermo Scientific™ TMTpro™ 18plex (cat. no. A52045) labeling was carried out as described by the manufacturer. After labeling completion, the samples were pooled and the peptides were desalted using 100 mg C18-SPR (waters) as described by the manufacturer protocol. The sample was lyophilized and phospho-peptides were enriched using High-Select Fe-NTA Phosphopeptide Enrichment (cat. no. A32992, Thermo Fisher Scientific). The enriched phosphopeptide fraction was then further fractionated using Pierce™ High pH Reversed-Phase Peptide Fractionation Kit (Pierce™ High pH Reversed-Phase Peptide Fractionation Kit, cat. no. 84868) was used. Fractionation protocol as described by the manufacturer kit. Eight fractions were generated from this step and were analyzed as follows:

#### LC-MS-MS

1 µg of each High pH Reversed-Phase enriched phosphopeptides was analyzed by ultra-high-pressure liquid chromatography (UPLC) coupled with tandem mass spectroscopy (LC-MS/MS) using nano-spray ionization. The FAIMS nano-spray ionization experiments were performed using an Orbitrap fusion Lumos hybrid mass spectrometer (Thermo Fisher Scientific) interfaced with nano-scale reversed-phase UPLC (Thermo Dionex UltiMate™ 3000 RSLC nano System) using a 25cm, 75 µm ID glass capillary packed with 1.7 µm C18 (130) BEH^TM^ beads (Waters corporation). Peptides were eluted from the C18 column into the mass spectrometer using a linear gradient (5–80%) of ACN (Acetonitrile) at a flow rate of 375 μl/minute for 120 minutes. The buffers used to create the ACN gradient were: Buffer A (98% H_2_O, 2% ACN, 0.1% formic acid) and Buffer B (100% ACN, 0.1% formic acid). Mass spectrometer parameters are as follows; FAIMS CV setting: -40, -60, and -80V and carrier nitrogen gas flow set at 3.8 liters per minute. An MS1 survey scan using the orbitrap detector (mass range (m/z): 400-1500 (using quadrupole isolation), 60000 resolution setting, spray voltage of 2200 V, Ion transfer tube temperature of 290°C, AGC target of 400000, and maximum injection time of 50 ms) was followed by data dependent scans (top speed for most intense ions, with charge state set to only include +2-5 ions, and 5 second exclusion time, while selecting ions with minimal intensities of 50000 at in which the collision event was carried out in the high energy collision cell (HCD Collision Energy of 38%) and the first quadrupole isolation window was set at 0.8 (m/z). The fragment masses were analyzed in the orbitrap detector (mass range (m/z): automatic scan with first scan at m/z= 100). The resolution was set at 30000 resolutions. AGC Target set to 30000, and maximum injection time: 54 ms. Protein identification and quantification was carried out using Peaks Studio X (Bioinformatics solutions Inc.). Global Proteome analysis of TMT labelled sample: The flowthrough peptides from the phospho-peptide enrichment step were fractionated using Pierce™ High pH Reversed-Phase Peptide Fractionation Kit (Pierce™ High pH Reversed-Phase Peptide Fractionation Kit, cat. no. 84868). Fractionation protocol as described by the manufacturer kit.

Eight collected fractions were each analyzed by ultra-high-pressure liquid chromatography (UPLC) coupled with tandem mass spectroscopy (LC-MS/MS) using nano-spray ionization. The FAIMS nano-spray ionization experiments were performed using an Orbitrap fusion Lumos hybrid mass spectrometer (Thermo Fisher Scientific) interfaced with nano-scale reversed-phase UPLC (Thermo Dionex UltiMate™ 3000 RSLC nano System) using a 25 cm, 75 µm ID glass capillary packed with 1.7 µm C18 (130) BEH^TM^ beads (Waters corporation). Peptides were eluted from the C18 column into the mass spectrometer using a linear gradient (5–80%) of ACN (Acetonitrile) at a flow rate of 375 μl/minute for 120 minutes. The buffers used to create the ACN gradient were: Buffer A (98% H_2_O, 2% ACN, 0.1% formic acid) and Buffer B (100% ACN, 0.1% formic acid). Mass spectrometer parameters are as follows: FAIMS CV setting: -40, -60, and -80V and carrier nitrogen gas flow set at 3.8 liters per minute. A MS1 survey scan using the orbitrap detector (mass range (m/z): 400-1500 (using quadrupole isolation), 60000 resolution setting, spray voltage of 2200 V, Ion transfer tube temperature of 290°C, AGC target of 400000, and maximum injection time of 50 ms) was followed by data dependent scans (top speed for most intense ions, with charge state set to only include +2-5 ions, and 5 second exclusion time, while selecting ions with minimal intensities of 50000 at in which the collision event was carried out in the high energy collision cell (HCD Collision Energy of 38%) and the first quadrupole isolation window was set at 0.8 (m/z). The fragment masses were analyzed in the orbitrap detector (mass range (m/z): automatic scan with first scan at m/z = 100. The resolution was set at 30000 resolution. AGC Target set to 30000, and maximum injection time: 54 ms. Protein identification and quantification was carried out using Peaks Studio X (Bioinformatics solutions Inc.).

#### Analysis

Identified phosphopeptides were normalized to their respective protein by dividing the intensity of the phosphopeptide by the intensity of its corresponding protein per sample. Phosphopeptides were removed if the intensity of the phosphopeptide or the intensity of its corresponding protein were below detection in a sample in a given comparison. Comparisons were made between WT females vs HET females, WT females vs HOM females, WT males vs HET males, WT males vs HOM males, HET females vs HET males, and HOM females vs HOM males. Significance was determined by student’s *t*-test and fold change was calculated from the normalized mean intensity. A p-value of 0.05 was selected to identify global trends in pathway alteration. Volcano plots show log-transformed p-values vs the log-transformed fold change quantified per change in normalized phosphopeptide abundance between comparisons. Next, Ontology enrichment analysis was conducted using EnrichR package (R package version 3.2, 2023)(62-64). Graphs display the top ten most significant ontologies for GO Biological Process 2023, GO Cellular Component 2023, and GO Molecular Function 2023 from the Gene Ontology Consortium(104, 105) (p<0.05). Additionally, heatmaps were generated to display phosphopeptide changes that trend (increase or decrease) with increasing ΔF48 alleles. These were selected by identifying normalized phosphopeptides significantly altered in HOM compared to WT with a corresponding trend (or significant change) in HET compared to WT for each sex. Heatmaps were also created for phosphopeptide changes that trend (increase or decrease) similarly in both male and female mice in WT vs HET or WT vs HOM. These similarly change phosphopeptides are defined as having the same trend in males and females with significant differences in normalized phosphopeptide abundance between WT vs HET or WT vs HOM in at least one sex. Analysis, statistics, and graphical representations were all performed in R programming (R version 4.3.3; R studio Version 2023.12.1+402)(106).

### Statistical Analysis

Statistical comparisons were made using one-way analysis of variance (ANOVA), Student’s *t*-test, or *χ*^2^-test where indicated. Sex was considered a biological variable and all tests utilized male and female mice. No significant differences were observed based on the age of mice. Outliers were determined using Grubbs’ test (α=0.05) where appropriate. All bar plots are shown as mean±S.E.M. Analysis, statistics, and graphical representations were prepared in R programming (R version 4.3.3; R studio Version 2023.12.1+402)(106).

### Study Approval

All procedures were approved by the Institutional Animal Care and Use Committee at the University of California, San Diego and were performed in accordance with the National Institutes of Health’s Guide for the Care and Use of Laboratory Animals, published by the National Academies Press (US), 2011, 8^th^ Edition.

## Data Availability

Phosphoproteomic data generated for this manuscript has been deposited onto the Proteomics Identification Database (PRIDE).

## Acknowledgements

We thank members of the Newton laboratory for their many helpful discussions and edits to the manuscript. We thank Ivan Florentino for technical support. We thank Dr. Joshua Mayfield for helpful discussions. We thank Dr. Susan Ackerman and laboratory for advice and helpful discussions.

## Author Contributions

A.C.N., S.S.T., and S.A.W. conceived the project, designed experiments, and wrote the manuscript. S.A.W., Y.M., C.C. performed the experiments. G.G. analyzed patient data and provided subject matter expertise. M.G. performed MS analysis and provided subject matter expertise. A.J.R. and S.W. performed behavioral experiments. A.J.R. provided subject matter expertise. C.A.P. and S.R.L. coordinated and generated the mouse model and provided subject matter expertise. S.A.W. analyzed data for and generated all figures. All authors edited the manuscript.

