## Supplemental Figures for "Sex specific disruptions in Protein Kinase Cγ signaling in a mouse model of Spinocerebellar Ataxia Type 14"

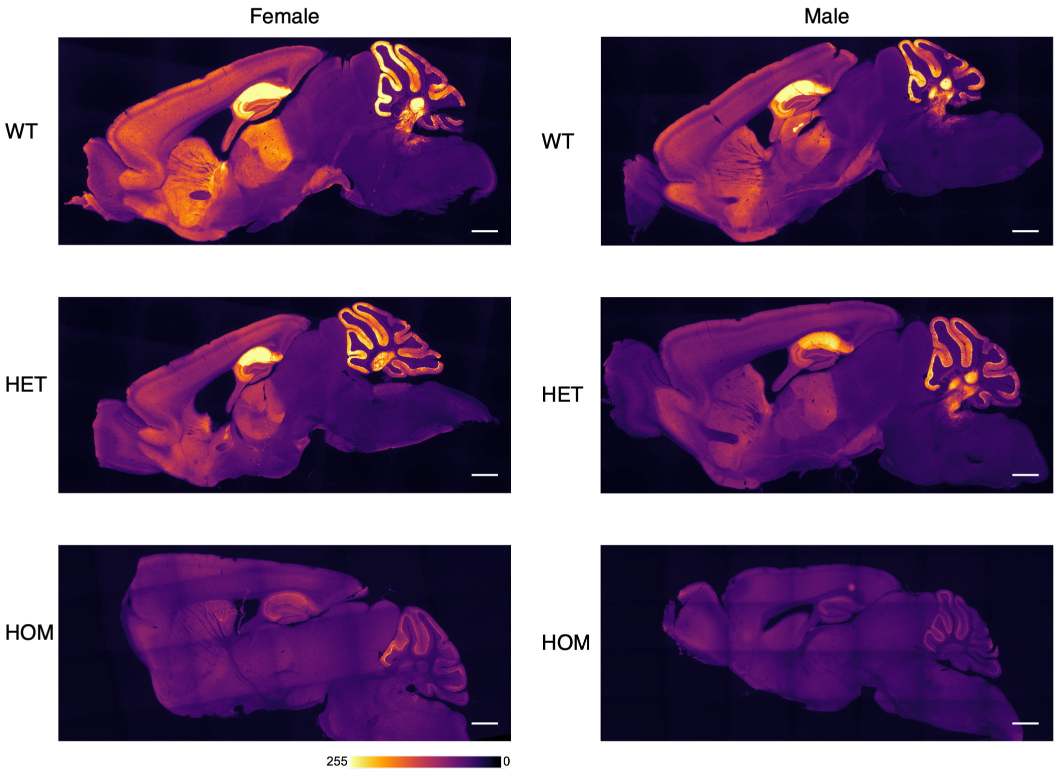


**Supplemental Figure 1: PKCγ is reduced in whole brain of SCA14 mice.**
Fluorescence immunohistochemistry of PKCγ was performed on fixed sagittal brain sections (40µm) from SCA14 mice of all genotypes (WT, HET, HOM) and sexes, and imaged using widefield microscopy on a Keyence Automate Imaging System microscope (Keyence Corporation). Representative images of PKCγ staining indicate a gene dose dependent loss of PKCγ in whole brain (Estimated scale bar=1mm, respective color scale indicates intensity).

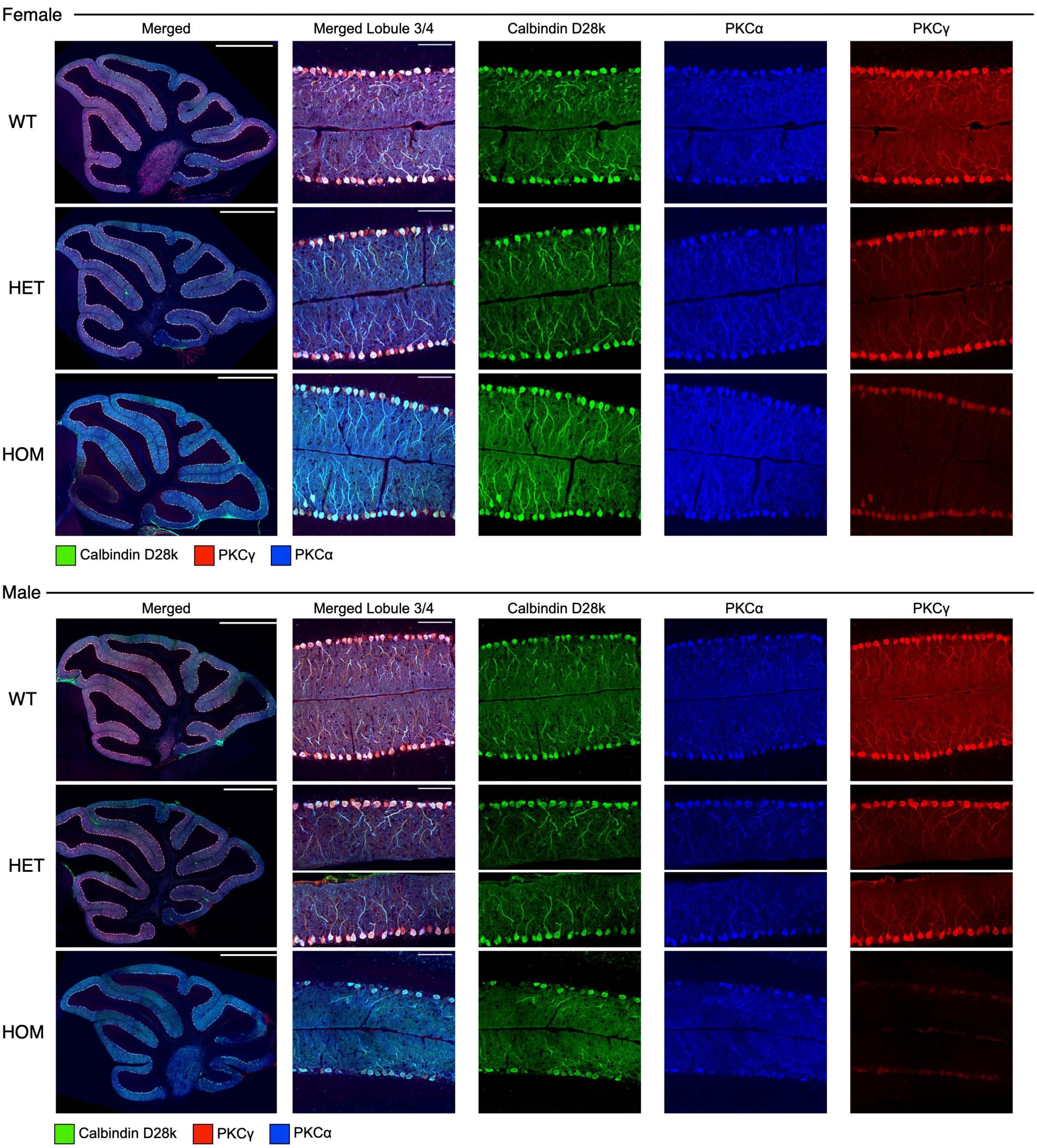


**Supplemental Figure 2: PKCγ and PKCα are localized to Purkinje cells of the Cerebellum.** Fluorescence immunohistochemistry was performed on fixed sagittal brain sections from all genotypes (WT, HET, HOM) and sexes. Shown are unquantified max projected images to allow visualization of the cerebellum and colocalization of Calbindin D28k (green), PKCγ (red), and PKCα (blue) in the whole cerebellum and a magnification of the PCL and ML in lobule 3 and 4 as a merged image as well as for each protein individually. Whole cerebellum scale bar=1mm, PCL and ML magnification scale bar=100μm.

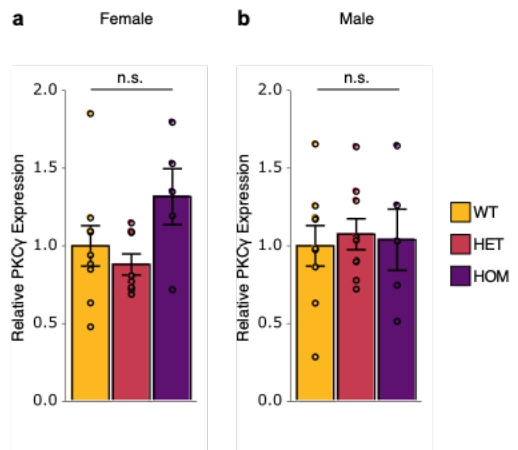


**Supplemental Figure 3: SCA14 mice have equivalent levels of PKCγ RNA in the cerebellum.** RT-qPCR was performed on RNA isolated from whole cerebellar homogenate from all genotypes (WT, yellow; HET, red; HOM, purple) and sexes to assess transcriptional changes. No significant changes in PKCγ RNA abundance were detected in **a)** female or **b)** male mice. The ΔΔCt method was used to quantify relative abundance and data is normalized to WT control group (N=5-9 mice per group). Bar graphs represented quantification of mean±S.E.M. Significance was determined by One-way ANOVA with Tukey's post hoc (not significant indicated by n.s.).

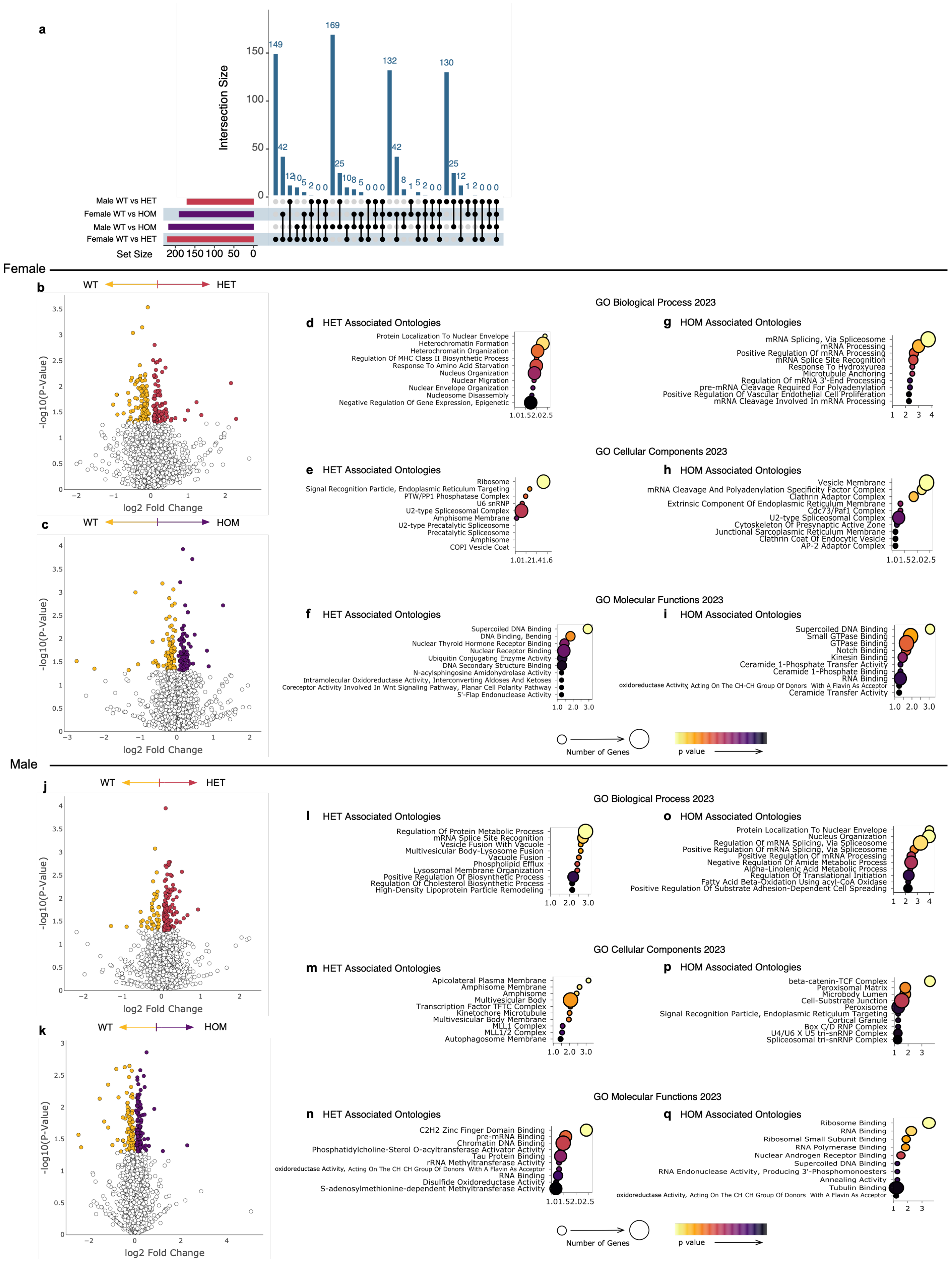


**Supplemental Figure 4: ΔF48 PKCγ alters the cerebellar proteome in SCA14 mice.**

Proteomic analysis was carried out on protein extracted from whole cerebellar homogenate from all genotypes (WT, yellow; HET, red; HOM, purple) and sexes (N=3 mice per group). A total of 5216 quantifiable proteins were detected. **a)** An upset plot summarizes the quantity of significantly different proteins and shared proteins identified between groups, with male HOM mice exhibiting the largest difference in the proteome compared to WT mice ($p<0.05$). Volcano plots show log-transformed p-values vs the log-transformed fold change quantified per change in protein abundance between WT compared to **b)** HET and **c)** HOM in female mice. Color represents phosphopeptides with p-value < 0.05. Dot plots indicate the top 10 most significantly enriched for gene ontologies from the differentially abundant proteins identified in **d-f)** WT vs HET and **g-i)** WT vs HOM females for biological processes, cellular components and molecular functions. Volcano plots show log-transformed p-values vs the log-transformed fold change quantified per change in protein abundance between WT compared to **j)** HET and **k)** HOM in male mice. Color represents phosphopeptides with p-value < 0.05. Dot plots indicate the top 10 most significantly enriched for gene ontologies from the differentially abundant proteins identified in **l-n)** WT vs HET and **o-q)** WT vs HOM males for biological processes, cellular components and molecular functions. Dot plots show ontology vs -log10(p-value), and plot color scale indicates p-value while dot size indicates number of genes per ontology ($p<0.05$).

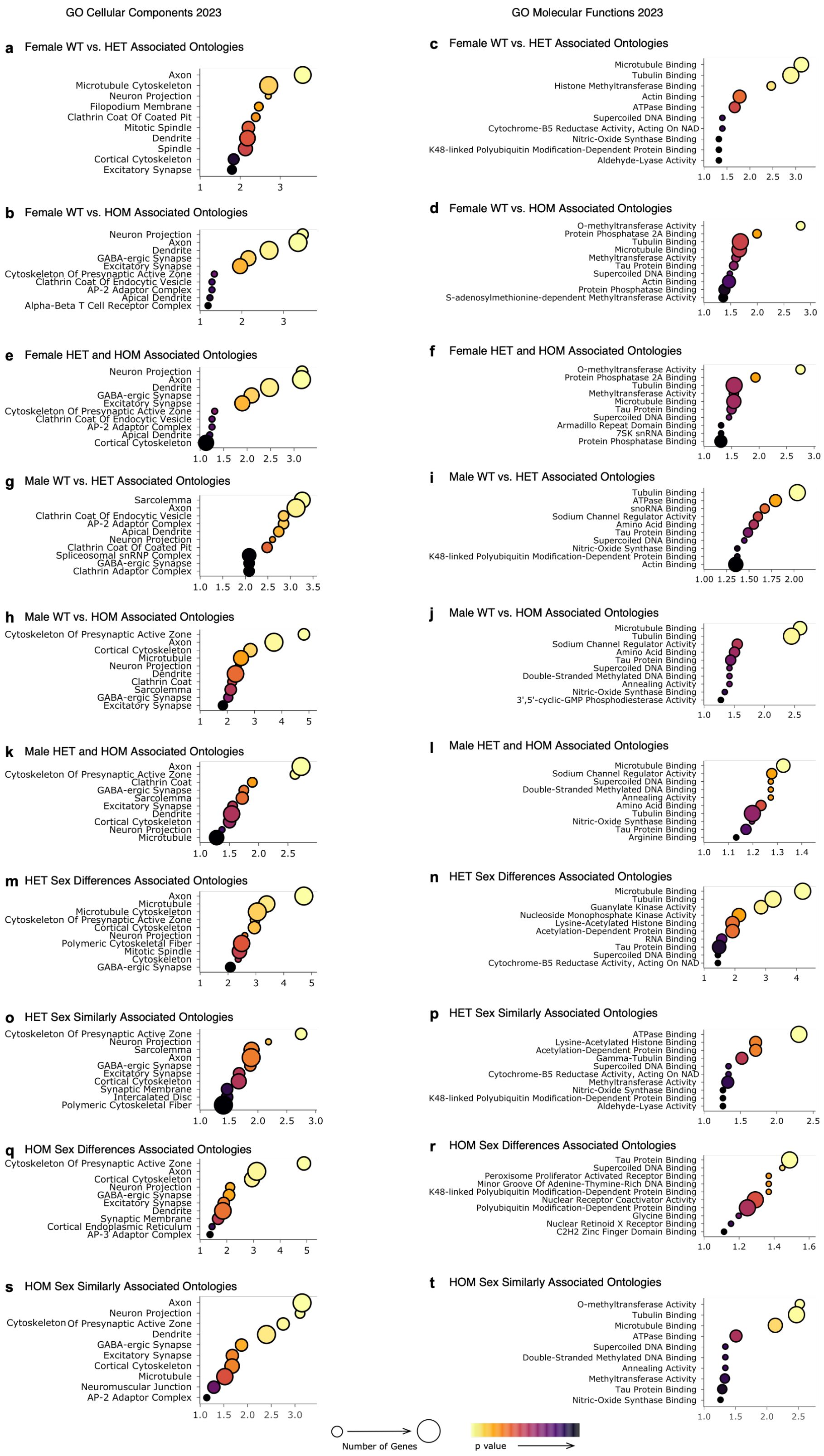


**Supplemental Figure 5: Ontology analysis of the SCA14 mouse phosphoproteome.**

Phosphoproteomic analysis was carried out on protein extracted from whole cerebellar homogenate from all genotypes (WT, HET, HOM) and sexes (N=3 mice per group). Changes in phosphopeptide intensity were normalized to the corresponding protein intensity. Significant differences in phosphopeptides abundance (as determined by two sample t-test <0.05) were compared between WT vs HET and WT vs HOM in female and male mice, or male vs female in HET and HOM mice. Gene ontology analysis was performed, and graphs indicate the top ten most significant ontologies for Cellular components (left) and Molecular functions (right) determined with EnrichR. Dot plots show ontology vs -log10(p-value), and plot color scale indicates p-value while dot size indicates number of genes per ontology ($p<0.05$).
